# Behavioral Signatures of Post-Decisional Attention in Preferential Choice

**DOI:** 10.64898/2026.01.10.698805

**Authors:** Ariel Zylberberg, Ian Krajbich, Michael N. Shadlen

## Abstract

Attention plays a key role in decision-making by directing limited cognitive resources to relevant information. It has been proposed that attention also biases the decision process, due to a multiplicative interaction between attention and subjective value (e.g., Krajbich et al., 2010). We tested two predictions of models that posit a causal multiplicative effect of attention on decision formation: (i) the last fixation should be more informative about the choice when the overall value of the alternatives is high, and (ii) more attention should be directed to the chosen option when choices conflict with stated preferences than when they do not. Reanalyzing several datasets from a food-choice task, we found no evidence supporting these predictions. An alternative model where attention reflects choices after the decision has completed, explains key observations, including the last-fixation bias, the gaze-cascade effect and the effect of the overall value of the alternatives on response times. However, this model does not fully account for the association between dwell time and choice. We conclude that gaze behavior prior to the choice report likely reflects both decisional and post-decisional processes.

## Introduction

Attention plays a key role in decision making, enabling individuals to focus on relevant information while ignoring distractions. It has also been hypothesized that attention biases decision makers’ preferences and choices. During decisions involving spatially distributed stimuli, gaze progressively shifts toward the option ultimately chosen, a phenomenon termed the gaze cascade effect (Shimojo et al., 2003; Glaholt and Reingold, 2009). Additionally, decision-makers often select the option they fixate on last before reporting their choice (a “last-fixation” bias; Krajbich et al. 2010). Experimental manipulations of gaze, including spatial cues, variations in exposure duration, and salience control, have demonstrated that options receiving more attention are more likely to be chosen (Nittono and Wada, 2009; Bhatnagar and Orquin, 2022; Pleskac et al., 2023). As gaze reflects spatial attention, these findings support the idea that attention biases choice (Shimojo et al., 2003; Armel et al., 2008; Zajonc, 1968).

The attentional drift diffusion model (*aDDM*) formalizes this hypothesis in a way that makes it suitable to quantitatively explain choice and response time (RT) (Krajbich et al., 2010). The *aDDM* builds on the drift-diffusion model of decision making (*DDM*), in which decisions are made by accumulating noisy samples of momentary evidence over time (Ratcliff, 1978). In preference-based decisions, the momentary evidence depends on the subjective value difference between the options. The decision is thought to terminate when the accumulated evidence crosses an upper or lower bound, simultaneously resolving the choice and the time it took to make it. The *DDM* and some of its variants, such as random walk, race, and attractor models, have been successful in explaining both choice and RT in a range of perceptual and cognitive decisions in which no role for attention is assumed (Link, 1975; Ratcliff and McKoon, 2008; Usher and McClelland, 2001; Vickers, 1979; Wang, 2002).

The *aDDM* extends the *DDM* by including a role for attention in the decision process. It proposes that attention can change the subjective value of decision alternatives: specifically, the value of unattended options is discounted by a multiplicative factor. This framework assumes that attention exerts its influence **intra-decisionally**—that is, during the deliberation process, before a choice is made. Originally developed for preference-based decisions between two options, the *aDDM* has been extended to multi-alternative (Krajbich and Rangel, 2011; Thomas et al., 2021), perceptual (Smith and Krajbich, 2019; Tavares et al., 2017), purchase (Krajbich et al., 2012), and attribute-based decisions (Fisher, 2021; Yang and Krajbich, 2023). The model provides an explanation for the association between choice, RT, subjective value, and gaze allocation, including the apparent causal influence of gaze on choice (for a review see Krajbich, 2019).

Here, we examine two behavioral predictions of models that posit a multiplicative influence of attention on subjective value. These predictions concern how gaze allocation (i.e., time spent viewing each item) relates to the overall value of the alternatives and to whether choices are consistent with stated preferences. One prediction of the *aDDM* and related models is that gaze allocation at the end of the decision should be more predictive of the choice when the overall value of the alternatives is higher (Ting and Gluth, 2025). A second prediction is that the difference in dwell time between items should depend strongly on choice consistency—specifically, that the lower-valued item should be looked at for longer for choices that are inconsistent with stated preferences. We tested the predictions by reanalyzing data from several previously published value-based decision tasks (Krajbich et al., 2010; Smith and Krajbich, 2018; Chen and Krajbich, 2016; Gwinn and Krajbich, 2016; Folke et al., 2016; Sepulveda et al., 2020). Contrary to these predictions, the data show systematic deviations from the *aDDM* and related frameworks (Callaway et al., 2021; Jang et al., 2021). Instead, the results suggest that the association between gaze and choice is at least partly **post-decisional**, arising after a covert commitment to a decision but before the overt response is executed (Cavanagh et al., 2014; Westbrook et al., 2020).

## Results

We analyzed data from previously published studies that used the food-choice paradigm introduced by Krajbich et al. (2010). In this paradigm, hungry participants performed two stages. In the first stage, participants viewed snack food items one at a time and rated how much they would like to consume each one (Fig. 1A); most of the studies we re-analyze used a numerical liking scale (Krajbich et al., 2010; Smith and Krajbich, 2018; Chen and Krajbich, 2016; Gwinn and Krajbich, 2016), while others elicited subjective value through an incentive-compatible Becker–DeGroot–Marschak auction (Folke et al., 2016; Sepulveda et al., 2020). In the second stage, participants were presented with pairs of items and asked to choose which one they would prefer to consume at the end of the experiment (Fig. 1B). Throughout the choice stage, participants’ gaze was recorded, allowing identification of the moments when attention was directed to the left versus right item (Fig. 1C).

**Figure 1.**
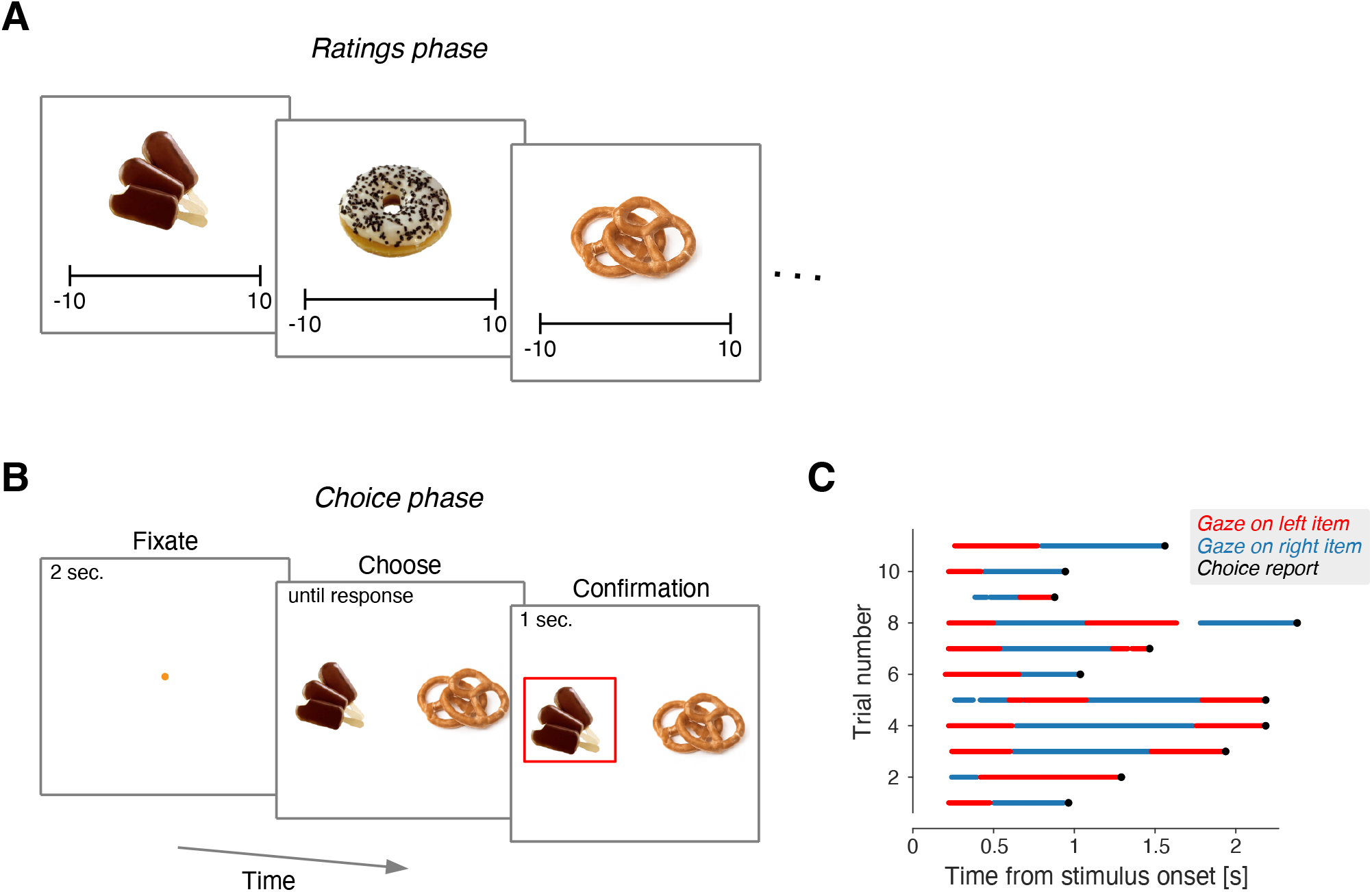
Food-choice task. The food-choice paradigm consists of two phases. In the first phase (A), participants were shown pictures of snack food items, one at a time, and were asked to rate how much they would like to consume each item, either on a numerical liking scale (Krajbich et al., 2010; Smith and Krajbich, 2018; Chen and Krajbich, 2016; Gwinn and Krajbich, 2016) or by stating their maximum willingness to pay in an incentive-compatible auction (Folke et al., 2016; Sepulveda et al., 2020). Note that while the original experiments used photographs of real packaged foods, the items depicted here are illustrative mockups. In the second phase (B), participants were presented with pairs of items and asked to choose which one they would prefer to consume at the end of the experiment. Participants responded manually; in most datasets, responses were made using the left and right arrow keys on a keyboard. (C) Gaze allocation between the left and right items shown from the moment both snacks appeared on the screen, for 11 representative trials from Krajbich et al. (2010). Red and blue indicate gaze directed to the left and right items, respectively. Times when the gaze was not directed to either item are left blank. Black dots indicate the time when the left or right key was pressed.

### The attentional drift-diffusion model

The decision-making process in the *aDDM* is governed by the state of a scalar decision variable, *x*, which takes the value zero at the start of each trial. The decision variable is updated according to

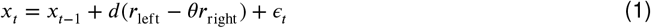

when the decision maker is looking at the left item, and according to

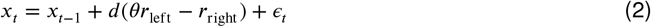

when looking at the right item. *r*_left_ and *r*_right_ represent the values assigned to the left and right items during the rating phase, respectively. If the decision variable reaches a bound at +1, the decision-making process terminates and the left item is selected, and if it reaches -1, the right item is selected. The parameter *d* controls the integration speed, *θ* is a parameter between 0 and 1 that determines how much the value of the unattended item is discounted, and *∈*_*t*_ is white Gaussian noise with variance σ^2^. The difference between the *aDDM* and the standard drift-diffusion model is that gaze modulates the drift rate through the parameter *θ*.

### The last-fixation bias on choice does not increase with overall value

A central feature of the *aDDM* is the multiplicative interaction between gaze and value. This multiplicative interaction accounts for faster response times when the overall value of the options is high—a phenomenon we refer to as the *magnitude effect on response times* (MERT) (Smith and Krajbich, 2019; Ratcliff et al., 2018). Given that attention has a greater influence when both options are highly valued, the last-fixation bias—the tendency to choose the item that was fixated last—should also be amplified under these conditions. A similar observation has recently been made by Ting and Gluth (2025). We refer to this prediction of the *aDDM* as the *magnitude effect on the last-fixation bias* (MELFB). We illustrate the MELFB prediction through simulations of the *aDDM* using the best-fitting parameters identified by Krajbich et al. (2010) and Smith and Krajbich (2019). To evaluate the impact of attention on choice, we apply logistic regression with the following model:

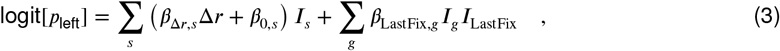

where Δ*r* represents the difference in value between the left and right items. *I*_*s*_ is an indicator variable that equals 1 for trials completed by subject *s* and 0 otherwise. *I*_LastFix_ is another indicator variable, set to 1 if the left item was fixated last and 0 if the right item was fixated last. We categorize trials into quintiles of *Σr*, the sum of the rating assigned to the left and right items; the variable *I*_*g*_ identifies trials belonging to quintile *g* ∈ {1..5}. The *β*s are the regression coefficients.

The first set of terms on the right-hand side of the equation (the summation over *s*) captures the effect of Δ*r* (along with a participant-specific bias) on the probability of choosing the left item. The second set of terms (the summation over *g*) reflects the influence of attention, measured by whether the left or right item was fixated last, with this effect estimated for each quintile of *Σr*.

Fig. 2A shows the regression coefficient *β*_LastFix,*g*_ for the different quintiles of overall value, for the simulations of the *aDDM* model. The dashed black line is not a fit to these data points, but is obtained from a model similar to that of Eq. 3 except that the term associated with the quantiles of overall value is replaced by an interaction between *I*_LastFix_ and *Σr* (Eq. 12). The analysis shows that the influence of the item attended last on choice increases with the overall value of the items (Fig. 2A).

**Figure 2.**
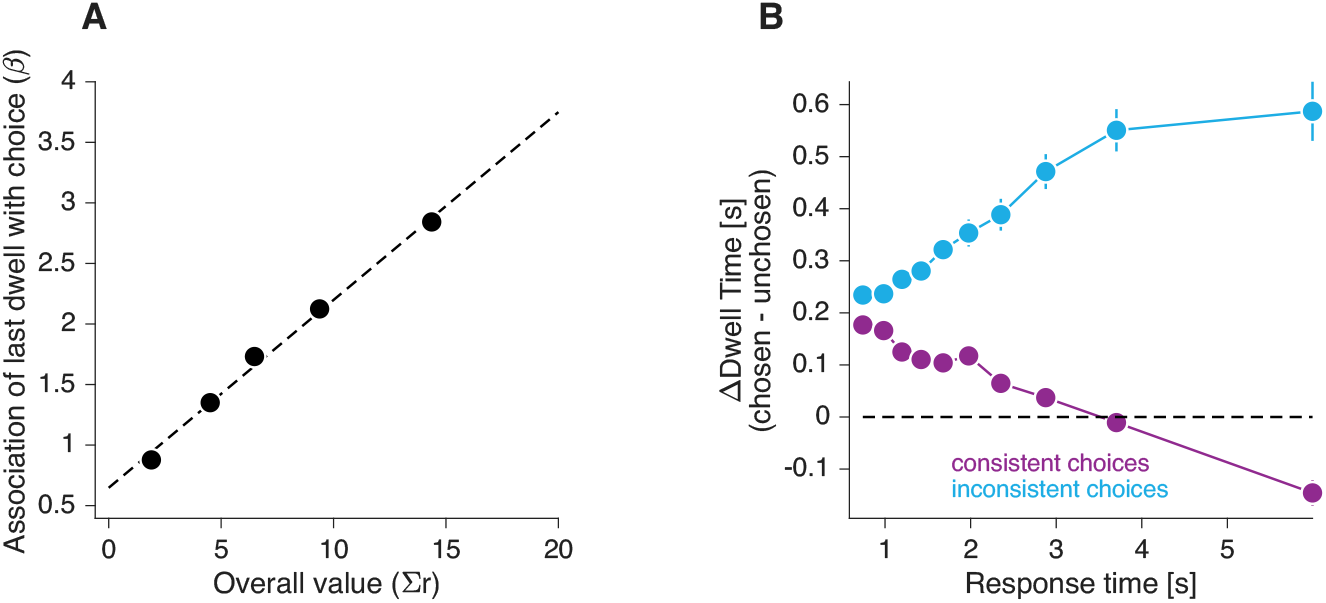
*aDDM* behavioral predictions. **(A)**Estimated strength of the association between last-dwell focus and choice (logistic regression coefficient; Eq. 3) as a function of overall value (Σ*r*). The five data points correspond to quintiles of the data, split by Σ*r*. The data were generated from simulations of the *aDDM* using the parameters reported in (Krajbich et al., 2010) and (Smith and Krajbich, 2019). The dashed line is derived from a related regression model that includes an interaction term between Σ*r* and whether the last fixation before the report was on the ultimately chosen or unchosen item. Error bars indicate s.e. **(B)**Time spent looking at the chosen item minus time spent looking at the unchosen item, as a function of response time. The data were generated from simulations of the *aDDM*. For each participant, trials were grouped into 20 categories defined by response-time decile and whether the choice was consistent with the initial ratings. Trials in which the two items received the same rating during the rating phase were excluded, because such choices cannot be classified as either consistent or inconsistent. The response times shown on the abscissa correspond to the mean response time across participants for each decile. Error bars indicate s.e. across participants.

### The difference in dwell time is independent of choice consistency

A second unexplored prediction of the *aDDM* is that the difference in total fixation time between the chosen and unchosen items, should depend on the consistency of the choice with the item ratings. We define inconsistent choices as those in which the lower-rated item is selected over the higher-rated one. The difference in total fixation time between the chosen and unchosen items is referred to as ΔDwell. According to the *aDDM*, if individual dwells are not influenced by item value, ΔDwell should be greater for inconsistent choices than for consistent ones. This prediction is confirmed in our simulations of the *aDDM* (Fig. 2B; p < 10^−7^, Eq. 13, *H*_0_: *β*_*c*_ = 0).

This prediction can be understood as follows. If, in a given trial, the decision maker had primarily attended to the higher-rated item, the likelihood of making an inconsistent choice would be lower than if attention had been equally divided between the options, because attention would have increased the extent to which the drift rate favored the higher-rated item. Thus, when a choice is known to be inconsistent, it is more likely that gaze was predominantly directed to the lower-rated item.

This logic also applies to consistent choices, but to a lesser extent. Even if attention is primarily allocated to the lower-valued item during a trial, it is still highly likely that the decision-maker will ultimately choose the higher-valued item. This is because, while attending to the lower-valued item reduces the effective drift rate in favor of the higher-valued item, it does not usually result in a change in the sign of the drift rate. Therefore, knowing that the decision was consistent does not provide as much information about the allocation of attention as knowing that the decision was inconsistent.

A similar logic explains why ΔDwell should depend on response time. The *aDDM* predicts that ΔDwell increases with RT for inconsistent choices (Fig. 2B). This is because the response time sets an upper limit on ΔDwell, so the longer the RT, the greater the value that ΔDwell can be. In contrast, ΔDwell decreases—and even becomes negative—with RT for consistent choices (Fig. 2B). This is because if the response time is longer than expected—given the values of the items being compared—then it is likely that attention was mostly focused on the lower-valued item. The effect is not as strong for inconsistent choices, because even if attention were primarily focused on the lower-valued item, the drift rate would still usually favor the unselected, higher-valued item.

### Testing the predictions in data from the food-choice task

We tested these predictions of the *aDDM* with data from the food-choice task. To test the prediction regarding the MELFB, we fit the logistic regression model (Eq. 3) to the data of Krajbich et al. (2010) and other datasets of the food choice task (Smith and Krajbich, 2018; Chen and Krajbich, 2016; Gwinn and Krajbich, 2016). In contrast with the prediction of the *aDDM*, the estimated strength of the association between last-dwell focus and choice was either non-significant or significantly negative (Fig. 3; p-values and *β*s indicated in the figure). A similar result was recently reported by Ting and Gluth (2025). That is, in contrast with model’s prediction, the last-fixation bias does not increase with the overall value of the alternatives.

**Figure 3.**
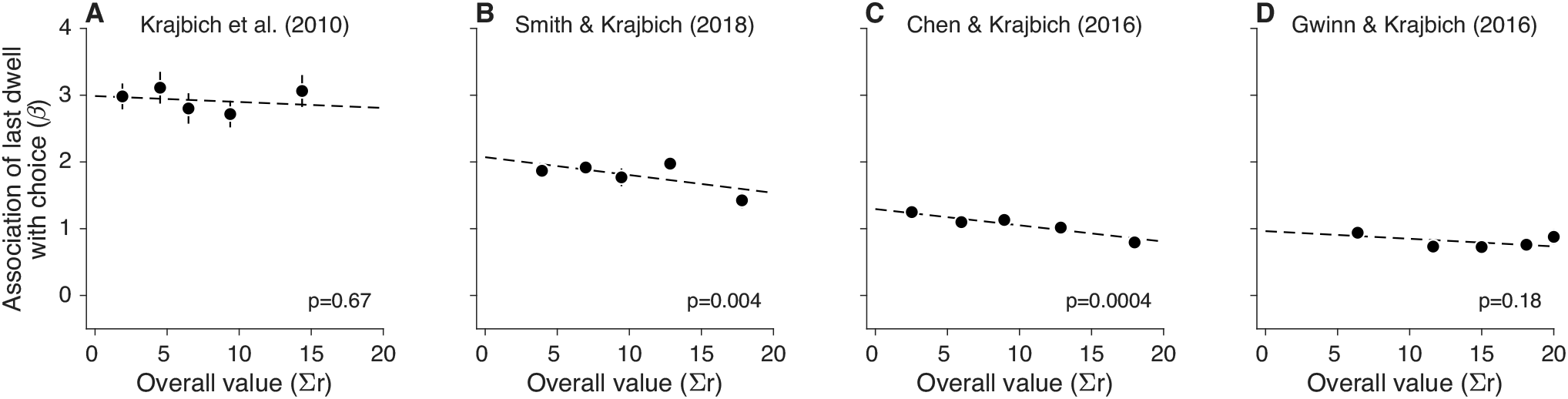
Association strength between last dwell and choice. Estimated strength of the association between last-dwell focus and choice as a function of Σ*r*. Same analysis and conventions as in Fig. 2A. Each panel shows data from a different dataset of the food-choice task: (A) Krajbich et al. (2010), (B) Smith and Krajbich (2018), (C) Chen and Krajbich (2016), (D) Gwinn and Krajbich (2016).

We also tested the prediction regarding the difference in ΔDwell between consistent and inconsistent choices. Across several datasets, we found no significant differences in ΔDwell for consistent and inconsistent choices (Fig. 4) (Eq. 13, *H*_0_ : *β*_*c*_ = 0; *p* > 0.13 for every dataset). This is incompatible with the prediction of the *aDDM*. This incompatibility is not specific to the *aDDM*, but as we show below, applies to other instantiations of models including those in which gaze is optimally allocated. [Note that some of these datasets were excluded from Fig. 3 because last-dwell focus was unavailable in the public data.]

**Figure 4.**
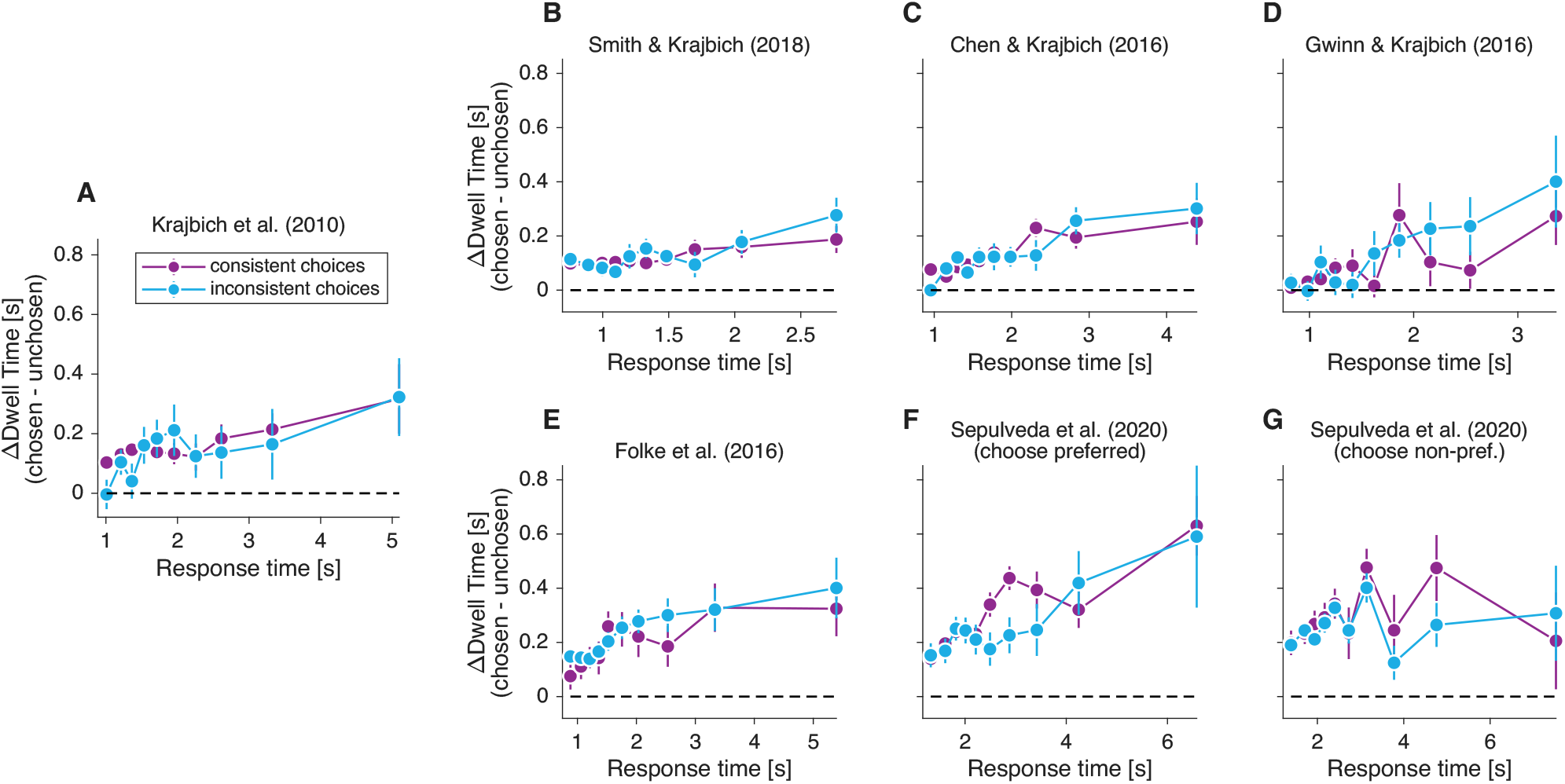
Difference in looking time for consistent and inconsistent choices. Time spent looking at the chosen item minus time spent looking at the unchosen item, as a function of response time. Same analysis and conventions as in Fig. 2. The seven panels correspond to behavioral data from (A) Krajbich et al. (2010), (B) Smith and Krajbich (2018), (C) Chen and Krajbich (2016), (D) Gwinn and Krajbich (2016), (E) Folke et al. (2016), (F-G) Sepulveda et al. (2020). In Sepulveda et al. (2020), participants either selected the item they preferred or the one they did *not* prefer. We analyzed the two variants separately.

### Is the gaze-choice association purely post-decisional?

The aspects of the data we analyzed in Figs. 3 and 4 are suggestive of a non-multiplicative interaction between attention and value (Cavanagh et al., 2014; Westbrook et al., 2020). We explore a model in which the link between gaze and choice arises only after a choice has been covertly made. Attention has no effect on the choice itself or on the time taken to make the choice. We refer to this model as the Post-Decision-Gaze (*PDG*) model.

In the *PDG* model, the decision is made by accumulating momentary evidence over time. The momentary evidence is represented by samples from a Gaussian distribution. The mean of the sampling distribution is a linear function of Δ*r*, such that the drift rate of the drift-diffusion process (*μ*) is given by:

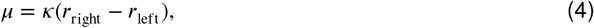

where *κ* is a signal-to-noise parameter. Unlike the *aDDM*, the drift-rate is not modulated by the focus of attention.

Unlike most implementations of the DDM, in the *PDG* model the variance of the momentary evidence depends on *Σr*. This assumption is needed to explain why response times depend on both Δ*r* and *Σr* (Ratcliff et al., 2018). We parameterize the variance as

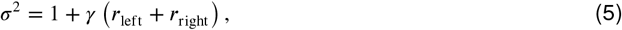

where *γ* is fit to the data. The assumption has empirical support in neurobiology (Discussion).

The evidence accumulation process begins after a short sensory delay, *τ*_s_ (Fig. 5A). For the purposes of our analysis, we fixed *τ*_s_ at 0.3 s for all participants. Note that while neurophysiological studies in monkeys have estimated that *τ*_s_ is on the order of 0.2 s (Roitman and Shadlen, 2002; Steinemann et al., 2022), *τ*_s_ is likely to be longer in the food-choice task since participants start each trial fixating on a central spot before directing their gaze to one of the choice alternatives (Krajbich et al., 2010).

**Figure 5.**
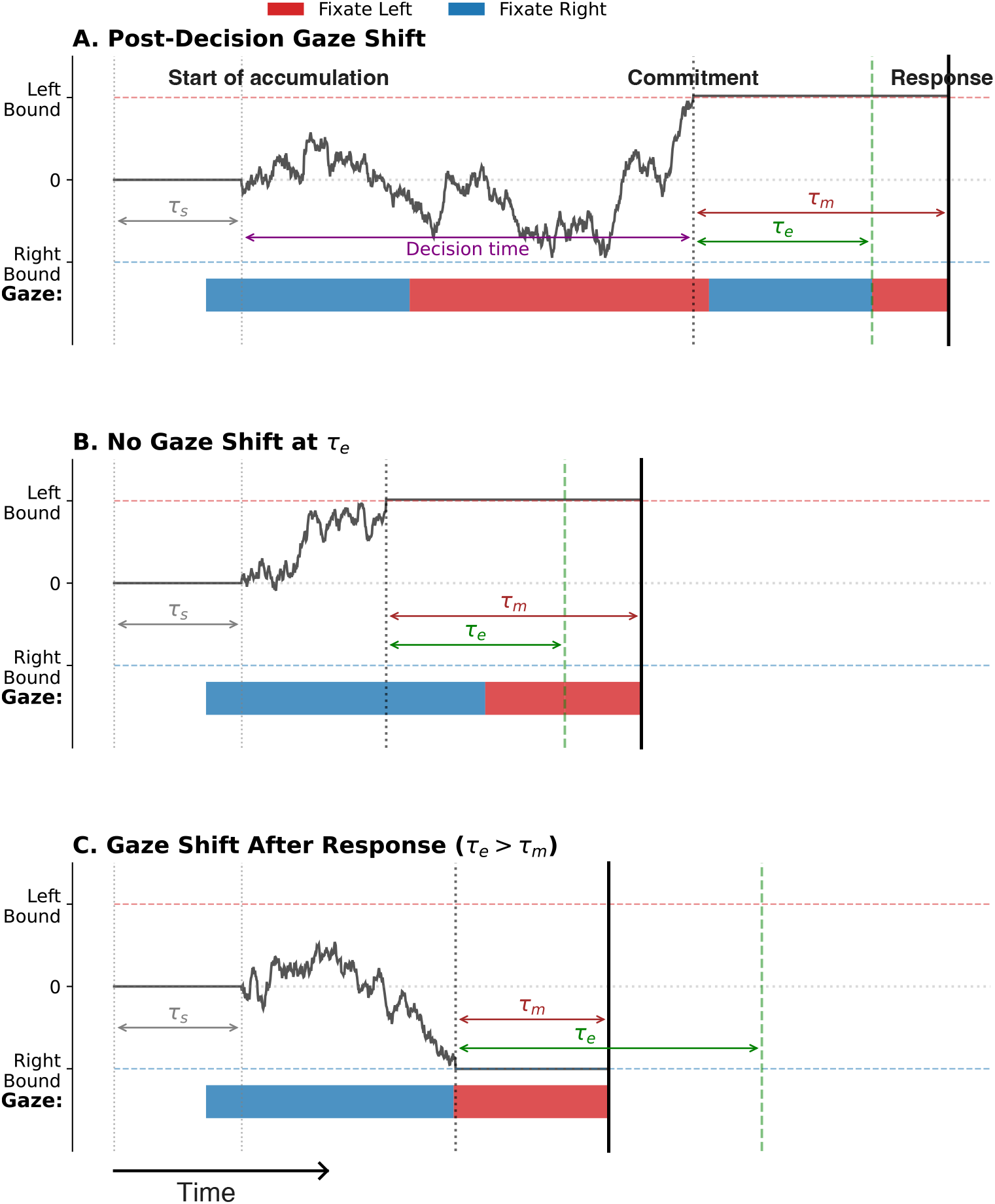
Sketch of the *PDG* model. (A) The decision is generated by accumulating momentary evidence over time until the process reaches either an upper or lower bound. Evidence accumulation begins after a sensory delay, τ_s_. Once a bound is crossed, an additional motor delay, τ_m_, elapses before the manual response is executed. Thus, the response time equals the decision time plus the non-decision delays τ_s_ and τ_m_. Gaze does not affect the decision process. Instead, at a random latency τ_e_ after the bound crossing, the gaze is directed toward the chosen option and remains there until the manual response. Because τ_e_ is typically shorter than τ_m_, the chosen item is usually the last fixated item before the response. (B) Example simulation in which the gaze is already on the chosen item at time τ_e_ following bound crossing; therefore, no gaze shift occurs. (C) Simulation in which the gaze shift to the chosen item occurs only after the manual response, and therefore does not influence pre-response gaze behavior. This explains why, in some trials—including the example shown—the non-chosen item is the last one fixated before the response.

The model also includes a non-decision latency, *τ*_m_, between the time when the decision maker commits to a choice, signaled by crossing the decision threshold, and the time when a key is pressed to report the choice (Fig. 5A). *τ*_m_ accounts for latencies related to motor preparation and is independent of decision difficulty. The response time is given by the sum of *τ*_m_, *τ*_s_, and the decision time (Fig. 5A).

In the model, attention has no causal effect on the decision process. Therefore, there is a 50% chance that the gaze will be directed to either item at the time a threshold is crossed. The key assumption of the model is that once the decision variable crosses the decision threshold, the gaze is directed to the chosen item. To account for eye-movement related latencies, we assume that directing gaze to the chosen item occurs with a latency of *τ*_e_ from the time of bound crossing (Fig. 5A), after which the gaze is held on the chosen item until the response. If the decision maker was already looking at the chosen item after *τ*_e_ has elapsed from the time of bound crossing, no additional gaze shift occurs (Fig. 5B).

Because the time it takes to make an eye movement is usually less than the time it takes to complete the manual response, *τ*_m_, the decision maker is more likely to be looking at the chosen item when a key is pressed (Fig. 5A,B). However, due to variability in *τ*_m_ and in *τ*_e_, the time at which gaze is directed to the chosen item after choice commitment varies from trial to trial, and may even occur after the key press, as in Fig. 5C. Crucially, in contrast with the *aDDM* and related models (Krajbich et al., 2010; Thomas et al., 2019; Krajbich and Rangel, 2011), it is the choice that affects gaze allocation, not the other way around.

### Fits of the *PDG* model to the behavioral data

We fit the *PDG* model to the choice and response time data from Krajbich et al. (2010). Fig. 6A and B show the proportion of trials in which participants chose the item on the right and the average RT as a function of Δ*r*. The *PDG* model provides a good fit to the choice and RT data.

**Figure 6.**
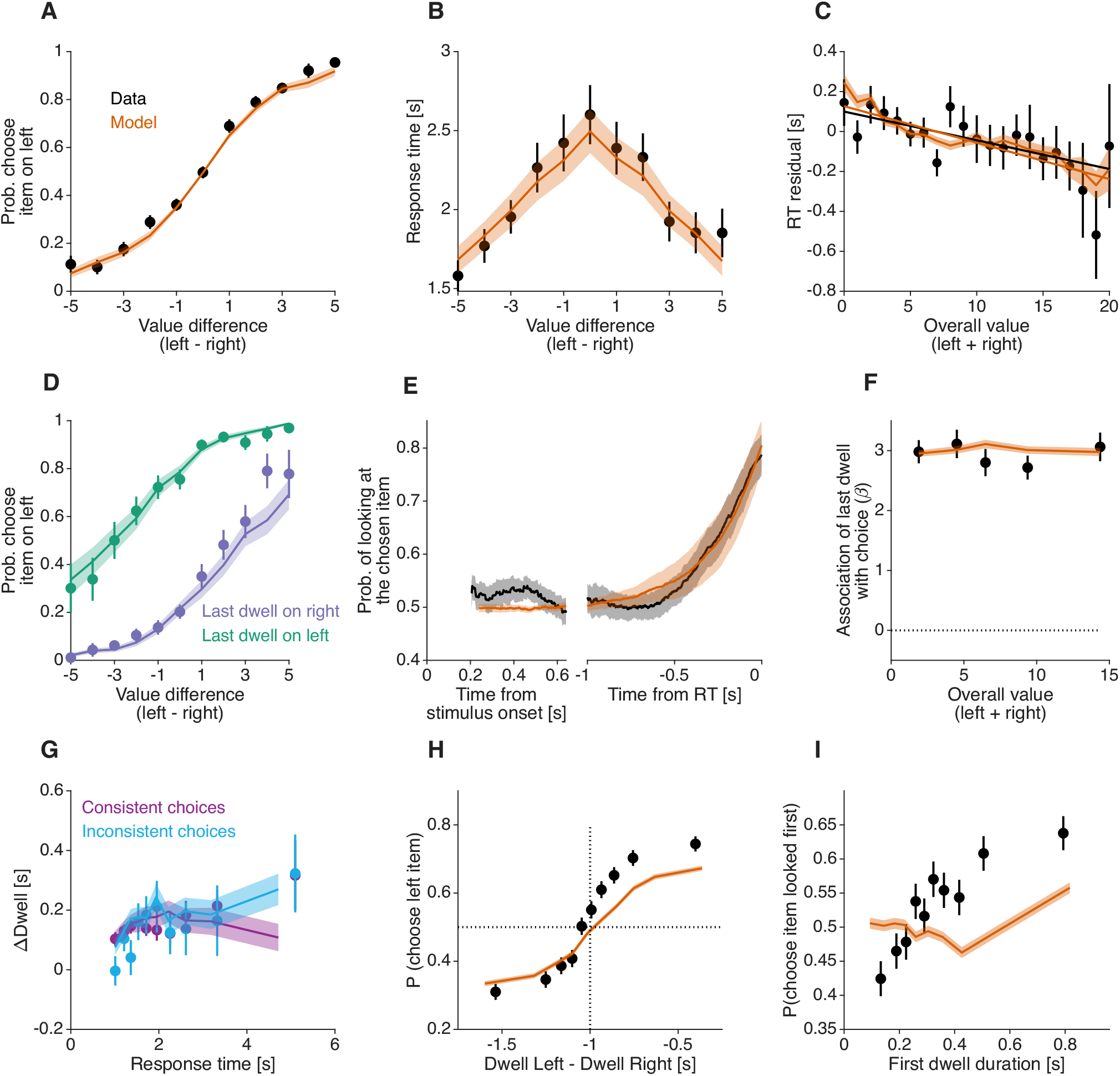
Data from Krajbich et al. (2010) and fits of the *PDG* model. (A) Proportion of trials in which the left item was selected as a function of the difference in value between the left and right items (Δ*r*). Points represent behavioral data and shading represents model fits. (B) Mean response time as a function of the value difference between the two items. (C) Residual response time (after subtracting the contribution of Δ*r*) as a function of the sum of the value of the two items presented in the trial (Σ*r*). Error bars indicate standard error of the mean (s.e.m.) across trials. (D) Proportion of trials in which the left item was selected, separated by whether the last fixation before the report was on the right (purple) or the left (green) item. (E) Probability that the decision maker is looking at the item that was ultimately chosen, plotted relative to the time since the two items were presented on the screen (left) and the response time (right). Error bands indicate 95% confidence intervals for the mean across participants. In the stimulus-aligned plot, the data are shown from the first moment that one of the two items is fixated on in at least 50% of the trials, which is ∼0.25 s. (F) MELFB for data (black) and model (orange). Same conventions as in Fig. 3. (G) Gaze bias for consistent and inconsistent choices. Same conventions as in Fig. 4. (H) Proportion of trials in which the left item was selected as a function of the difference in dwell time between the left and right items. Data (black) and model (orange) were grouped in deciles of the dwell difference, separately for each participant, and then averaged across participants. (I) Proportion of trials in which the item looked at first was selected, as a function of duration of the first dwell. Data (black) and model (orange) were grouped in deciles of the first dwell duration, separately for each participant, and then averaged across participants. In all panels except panel D and G, data are shown in black and model simulations are shown in orange. Error bars and error bands, unless otherwise noted, show the standard error of the mean (s.e.m.) across participants (N=39 for both model and data).

The model also accounts for MERT—the tendency to make faster decisions when the items being compared are overall more desirable, even when the value difference between them is the same (Smith and Krajbich, 2019; Sepulveda et al., 2020; Ratcliff et al., 2018). To illustrate this effect in Krajbich’s data, we fit a bell-shaped curve to the relation between RT and Δ*r*, and computed residuals of RT by subtracting from each trial’s RT the expectation given by the best-fitting bell-shaped curve (see Fig. S1 for an illustration of the method). We then analyzed how the residuals of RT correlated with *Σr*. This correlation was negative, indicating that responses were faster when *Σr* was higher. That is, the data show a magnitude effect on RT (Fig. 6C). Note that the analysis of RT residuals rather than raw RT is necessary because of the positive correlation between Δ*r* and *Σr* present in the data.

The *DDM* has been considered incapable of explaining the magnitude effect on RT because in the most common version of the *DDM* the drift rate depends only on the difference in value between options and ignores their absolute value, leading to the erroneous prediction that the response time for a given Δ*r* would be independent of *Σr*. The *PDG* model captures the magnitude effect because the variance of the momentary evidence is allowed to change with *Σr* (Eq. 5) (Ratcliff et al., 2018). For the best-fitting model, the variance of the momentary evidence increased with *Σr* (Table S1). Because higher variance leads to faster responses (Zylberberg et al., 2016), the model displays a magnitude effect on RT, similar to the data (Fig. 6C).

In the *PDG* model, attention has no causal influence on choice. Yet, it successfully accounts for several features of the observed association between gaze and choice. We compared the probability of choosing the right item—as a function of Δ*r* —on trials in which the last fixation was on the right versus the left item. The model predicts a systematic relation between the last fixation and choice that closely mirrors what is observed in the data (Fig. 6D).

In the *PDG* model, a gaze bias toward the chosen item can only occur during the non-decision time between choice commitment and report, giving rise to a *gaze cascade* effect—the observation that the probability of looking at the ultimately chosen item aligned to the response increases gradually over time (Fig. 6E). This gradual increase is explained as the average of step-like events (saccades to the chosen item) that occur at different times with respect to the response time. The variation in timing is due to trial-to-trial variability in non-decision time (captured by the model parameter σ_nd_, see Methods) and in *τ*_e_. Since the model was fit to maximize the likelihood of its parameters based on the choice and RT data, without using gaze information, the gaze cascade effect can be considered a prediction of the model.

The *PDG* model also correctly predicts the two novel behavioral observations shown in Fig. 3 and Fig. 4. In the *PDG* model, there is no significant change in the strength of the last-fixation bias as a function of the overall value of the alternatives, consistent with the experimental data (Fig. 6F). This is because, in the model, the probability of directing gaze to the covertly chosen item is independent of variables affecting the decision process, like Δ*r* or *Σr*.

The model also correctly predicts that ΔDwell does not depend on the consistency of the choice. In the model, ΔDwell is often positive because gaze is directed to the covertly chosen item and because *τ*_e_ is usually smaller than *τ*_m_. These factors do not depend on the consistency of the choice, resulting in similar values of ΔDwell for consistent and inconsistent choices (p = 0.27, Eq. 13, *H*_0_ : *β*_*c*_ = 0)(Fig. 6G).

### The *gaze cascade* effect continues after the choice report

In the *PDG* model, the bound crossing may have occurred hundreds of milliseconds before the choice is reported, and the two events are not time-locked due to variability in *τ*_m_ and *τ*_e_. The model posits that the gaze cascade arises because shifting gaze to the chosen item takes time, and the likelihood of having completed that shift increases with time elapsed since boundary crossing. By this logic, the model predicts an even further increase in the probability of looking at the chosen item immediately after the choice report.

To test this prediction, we examined the gaze allocation after the choice report. Fig. 7A shows the probability of looking at the chosen item, as a function of time, aligned to the response. The probability of looking at the chosen item continues to increase after the choice report. This can be seen more clearly in Fig. 7B, which shows the probability of looking at the chosen item during the 200 ms immediately before and after the choice report. For most participants, the probability is greater after the choice report (p < 10^−7^, Wilcoxon signed-rank test).

**Figure 7.**
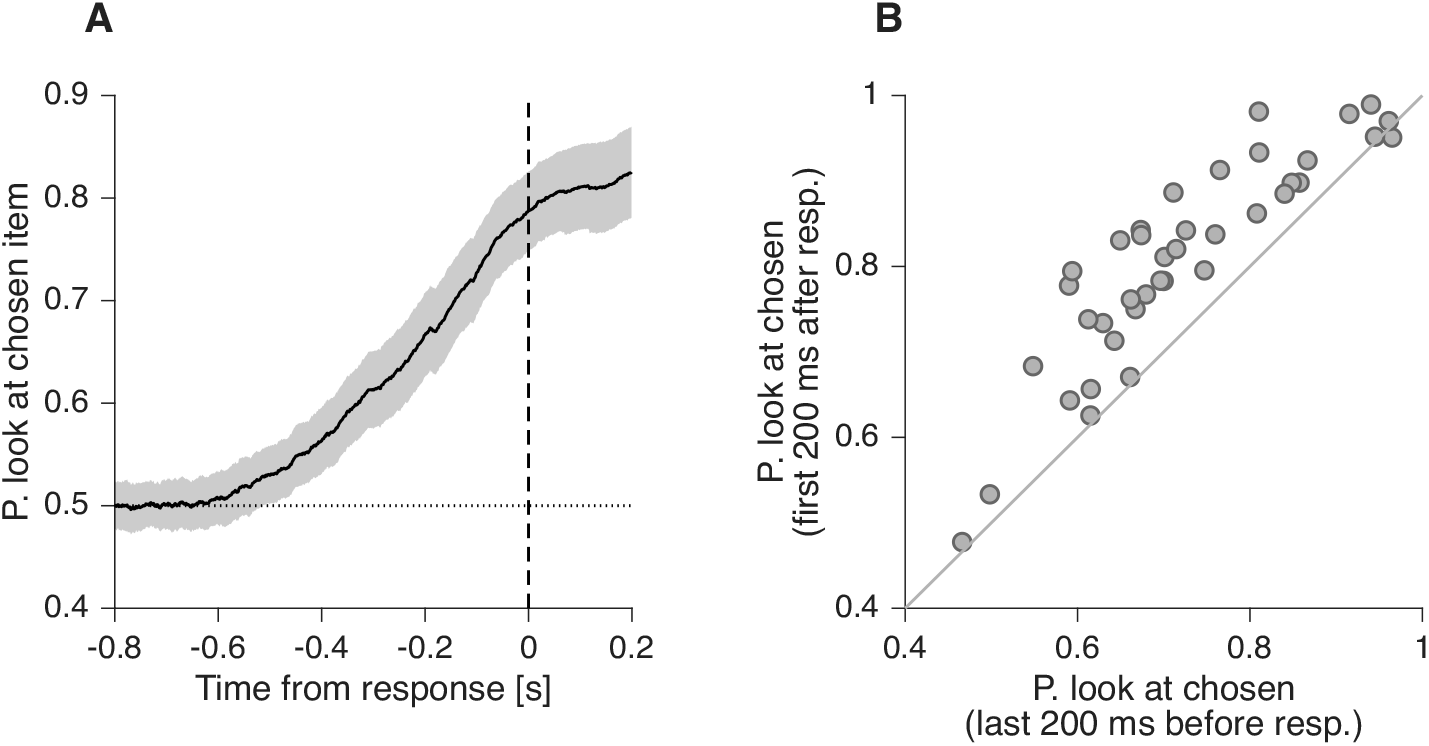
Gaze allocation after the choice report. (A) Probability that the decision maker is looking at the item that was ultimately chosen, aligned to RT. This probability increases even after the choice report. Error bands indicate 95% confidence intervals for the mean across participants. (B) Proportion of time that the decision maker is looking at the item that was ultimately chosen, calculated for the last 200 ms before the response (abscissa) and for the first 200 ms after the response (ordinate). Each data point represents one participant. Proportions were calculated as the sum of the time spent looking at the chosen item divided by the time spent looking at either one of the items (i.e., we exclude the times when the gaze was not directed at one of the two items).

That is, a common process—directing the gaze to the chosen item—may explain the allocation of gaze both immediately before and after the choice report. In contrast, models that posit an exclusively intra-decision effect of attention on choice must then explain the allocation of gaze after the response as a separate process (e.g., directing gaze to the chosen item), or by assuming that individuals continue to evaluate the decision alternatives even after the choice report.

### Parameter-sensitivity analysis

In the *PDG* model, the gaze–choice association arises from the temporal gap between manual and oculomotor response latencies, that is, from the difference *τ*_m_-*τ*_e_ (Fig. 6D–I). In our simulations, *τ*_e_ is modeled as a Normal distribution with mean *μ*_*e*_ = 0.35 s and standard deviation σ_*e*_ = *μ*_*e*_/3, truncated at zero to exclude negative values. To assess the sensitivity of the model to this assumption, we varied *μ*_*e*_ across a range and recomputed the key behavioral signatures. Fig. 8 shows the resulting gaze-cascade effect, the association between last-fixation focus and choice, and the interaction between ΔDwell and consistency, paralleling the analyses reported in Fig. 6E–G.

**Figure 8.**
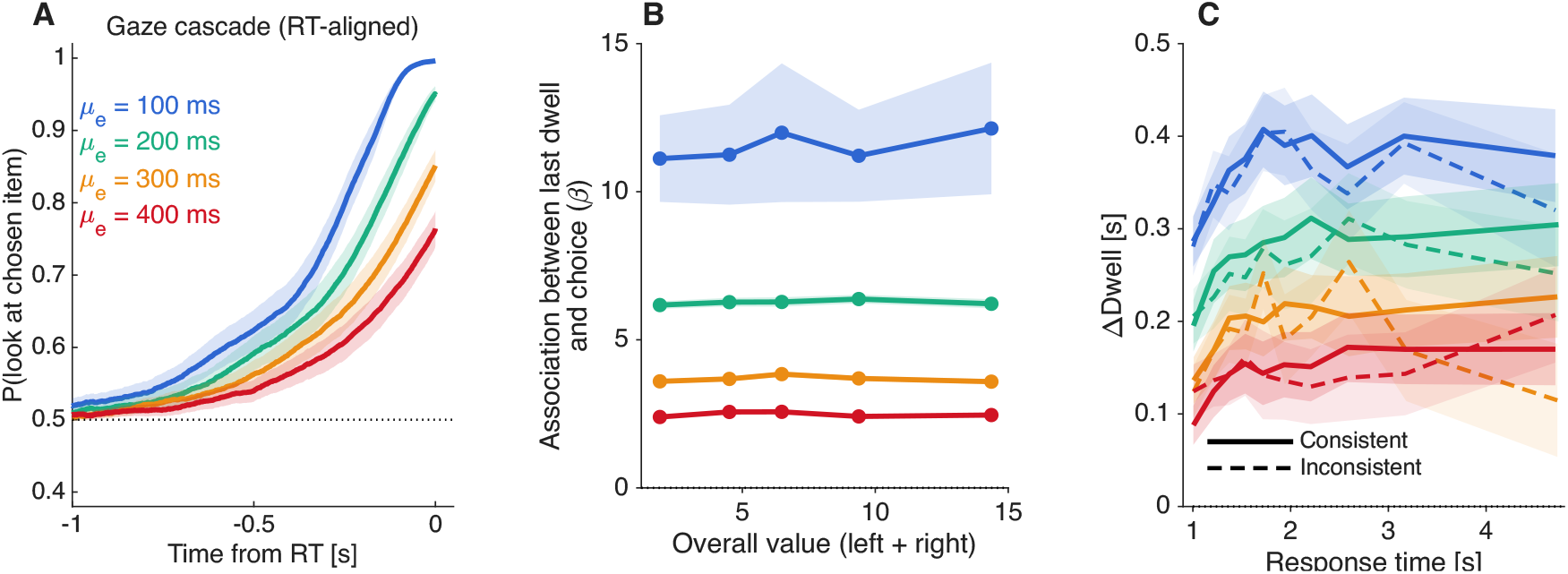
Sensitivity of the gaze–choice association in the *PDG* model. Simulations of the *PDG* model across a range of eye-movement latency values (*μ*_*e*_), assessing the robustness of the model’s predictions. **(A)** Predicted gaze-cascade effect. Similar analysis to that in Fig. 6E. **(B)** Predicted association between last-dwell focus and choice. Similar analysis to that in Fig. 6F. **(C)** Predicted ΔDwell (chosen minus unchosen) for consistent and inconsistent choices as a function of response time. Similar to the analysis in Fig. 6G.

As *μ*_*e*_ increases, the probability that gaze is already directed toward the chosen item at the time of response decreases (Fig. 8A). Because both *τ*_e_ and *τ*_m_ are variable, longer mean eye-movement latencies reduce the likelihood that the eye movement is completed before the manual response. Accordingly, the predicted association between gaze and choice weakens as *μ*_*e*_ increases (Fig. 8B). The magnitude of ΔDwell also declines with increasing *μ*_*e*_ (Fig. 8C). However, for all values of *μ*_*e*_, ΔDwell remains positive and does not differentiate between consistent and inconsistent choices. Thus, the qualitative predictions of the *PDG* model are robust to the precise value assumed for *τ*_e_. In the Discussion, we explain why the comparatively large best-fitting values of *τ*_e_ do not necessarily conflict with shorter post-decisional eye-movement latencies reported in neurophysiological studies of monkeys (Roitman and Shadlen, 2002).

### Limitations of the *PDG* model

Despite the ability of the *PDG* model to explain many aspects of the data, other aspects are not as well captured. In the *PDG* model, choice accuracy is unaffected by fixation patterns prior to crossing the decision threshold. That is, whether decision-makers focus more on the higher-value item or distribute their attention evenly between the options, the model’s predictions for choice and RT remain unchanged. This invariance arises from the lack of a causal relation between gaze and the decision-making process. As a result, the *PDG* model makes no concrete predictions for the gaze pattern before choice commitment.

In simulated data—where dwell durations are independent of the value of the alternatives—the *PDG* model fails to explain certain experimental findings that demonstrate an association between dwell duration and choice (Krajbich et al., 2010; Callaway et al., 2021). We highlight this limitation with two analyses. The first examines the likelihood of choosing the item on the right as a function of the difference in dwell time between the right and left items. Both empirical data and *PDG* model simulations reveal a positive association, but this relation is steeper in the data (Fig. 6H). That is, ΔDwell is more predictive of choice in the experimental data than in the model, suggesting that decision-makers may fixate longer on the eventually chosen item even before covert choice commitment.

The *PDG* model also fails to capture the relation between the duration of the first dwell and choice. Fig. 6I shows the proportion of trials in which decision-makers selected the initially fixated option as a function of the first dwell duration. This relation is steeper in the data than in the model. The *PDG* model does predict a positive relation between the duration of the first dwell and choice, but this is only because the bound is crossed during the first dwell on some trials. On trials with more than one dwell, the *PDG* model predicted no significant association between first-dwell duration and choice probability (all *p* > .05, likelihood-ratio tests; Eq. 14, Fig. 9A). In contrast, the empirical data revealed a significant positive association for shorter dwell-count conditions. Specifically, longer first-dwell durations significantly increased the probability of choosing the first-fixated item in 2-dwell (p = .013) and 3-dwell trials (p = .026), whereas this relationship was not significant for 4-dwell (p = .72) or 5-dwell trials (p = .96; Fig. 9B).

**Figure 9.**
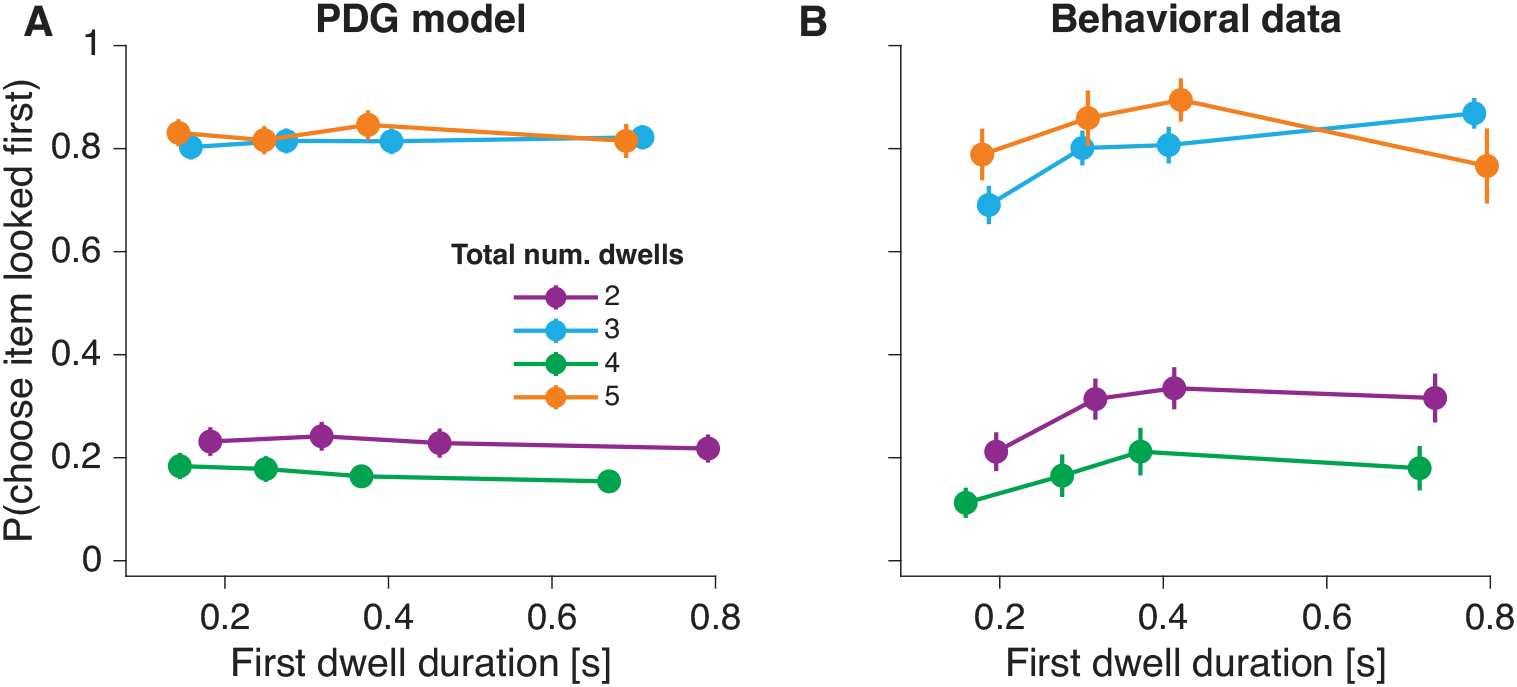
Association between first-dwell duration and choice probability. (A) Model-predicted probability of choosing the option that was fixated first, as a function of the duration of the first dwell. Trials are grouped by the total number of dwells (2–5), shown in separate colors. Data were binned into quartiles of first-dwell duration and then averaged across participants. Error bars indicate s.e.m. (B) Same analysis as A, for the behavioral data.

### Predictions of other decision making models

For comparison with the *PDG* model, we analyze different decision making models fit to the data of Krajbich et al. (2010). These models include variants of the aDDM (additive attention, inter-trial drift-rate variability, and post-decisional latencies) as well as pseudo-optimal models of decision making.

#### Additive intra-decision attention

We fit an additive variant of the *aDDM* to the data from Krajbich et al. (2010), using identical fitting procedures. In this additive intra-decisional attention model, the decision variable evolves according to

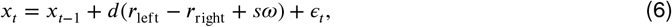

where *s* = +1 when attention is directed to the left item and *s* = −1 when it is directed to the right item. In this formulation, the drift rate shifts by +*ω* or −*ω* depending on which item is attended, in contrast to the *aDDM*, where the unattended item’s value is discounted multiplicatively.

The additive model, just as the multiplicative variant (Fig. 10A), fails to capture qualitative aspects of the data, most notably the interaction between choice consistency and ΔDwell (Fig. 10B).

**Figure 10.**
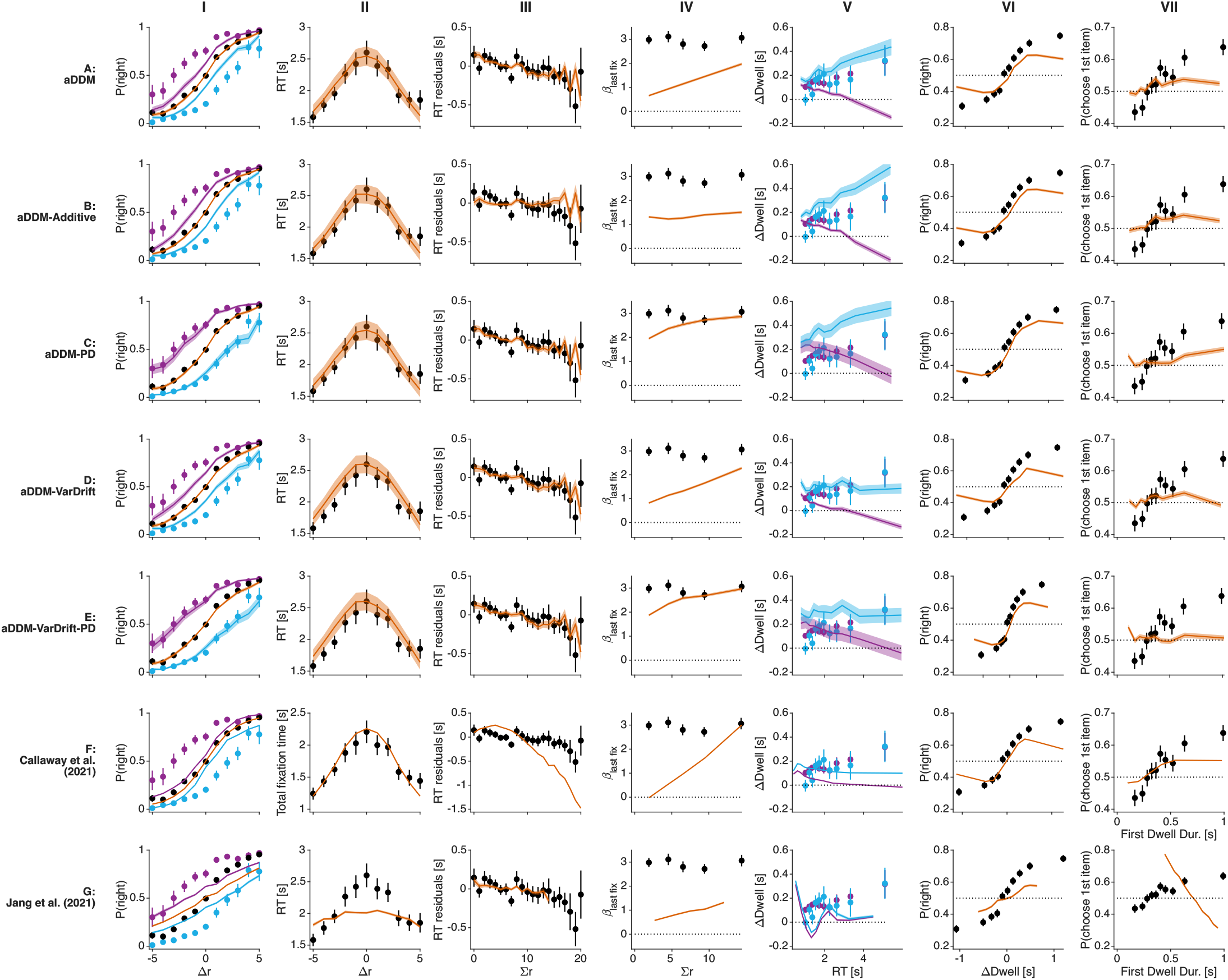
Predictions from different models. Fits of different models to the data of Krajbich et al. (2010). Same analyses and conventions as in Fig. 6. (A) Data and fits of the *aDDM* model. (B) *aDDM* with additive instead of multiplicative attention. (C) *aDDM* with post-decision attention. (D) *aDDM* with inter-trial variability in the drift-rates. (E) *aDDM* with inter-trial variability in the drift-rates and post-decision attention. (F) Model from Callaway et al. (2021). (G) Model from Jang et al. (2021).

#### Combined intra- and post-decision attention

Because neither a purely intra-decision nor a post-decision account of the gaze-choice association fully captures the behavioral data, we considered the possibility that a combination of the two accounts provides a better explanation (Westbrook et al., 2020). To this end, we combined elements from the *aDDM* and *PDG* models. In this variant of the model, prior to committing to a choice, decision dynamics follow the *aDDM* framework—the value of the unattended option is multiplicatively discounted and evidence accumulation terminates upon reaching a threshold at ±*B*. Following threshold crossing, the gaze shifts to the covertly chosen item after a delay *τ*_e_, as described in the *PDG* model.

Incorporating these post-decisional gaze shifts into the *aDDM* (Fig. 10C) improves the model’s ability to explain the dependency between the last dwell, overall value, and choice (compare Fig. 10A and Fig. 10C, columns I and IV). However, this hybrid approach still (*i*) inaccurately predicts that the gaze bias is stronger for inconsistent than for consistent choices (Fig. 10C, col. V), and (*ii*) displays a weaker association between the duration of the first fixation and choice, compared to what is observed in the data (Fig. 10C, col. VII). From this we conclude that the combination of intra- and post-decision attentional mechanisms does not fully account for the behavioral data.

#### Drift-rate variability across trials

None of the models that treat attention as an intra-decisional process were able to account for the difference in gaze bias between choices consistent versus inconsistent with stated preferences. We considered the possibility that inter-trial variability in the drift rate (e.g., Ratcliff and McKoon, 2008) could explain this discrepancy.

To test this, we fit a variant of the *aDDM* in which additive Gaussian noise corrupts the drift rate on each trial. The behavioral data and model fits are shown in Fig. 10D. The model still incorrectly predicts a larger gaze bias for inconsistent choices than for consistent choices. It also failed to capture the relation between first-dwell duration and choice.

To assess whether post-decisional processes might improve the model’s match to the data, we incorporated a post-decision gaze shift mechanism, similar to that in the *PDG* model. Even with this post-decisional mechanism the model predicts a larger gaze bias for inconsistent than for consistent choices, unlike what is observed in the data, and still fails to capture the observed association between first-dwell duration and choice (Fig. 10E).

We also explored an alternative implementation of inter-trial drift-rate variability, in which the item values— rather than the drift rate—are perturbed by additive Gaussian noise that remains constant within a trial. This approach is intended to capture the possibility that the values reported during the rating phase may differ from those used in the choice phase (c.f. Zylberberg et al., 2024). However, this variant did not provide a better fit to the behavioral data than the previous implementation (Fig. S4B,F-I).

#### Optimal gaze allocation

The allocation of attention in the *aDDM* is exogenous to the decision process—that is, attention shifts between items independently of the internal dynamics of decision making. In contrast, more recent studies model the control of attention as endogenous, arising from an optimization that balances the cost of delaying the decision and collecting additional evidence, with the expected benefit of improved accuracy (Callaway et al., 2021; Jang et al., 2021; Song et al., 2019; Hébert and Woodford, 2018; Gluth et al., 2026, 2020; Zhu, 2022). For instance, Callaway et al. (2021) formalized the decision process as a partially observable Markov decision process (POMDP), and approximated its solution by assuming that the value of additional deliberation can be expressed as a linear combination of factors such as the expected reward from acquiring an extra evidence sample and the expected reward if the item values were perfectly known.

We asked whether the optimal attentional allocation model of Callaway et al. (2021) would show the same discrepancies with the behavioral data as the *aDDM*. To this end, we reanalyzed the simulations of the optimal model performed by Callaway and colleagues. Like the *aDDM*, the model of Callaway et al. (2021) incorrectly predicts a strong effect of overall value on the strength of the association between attention and choice (Fig. 10F). This is clearly counter to what is observed in the data (Fig. 3).

We also analyzed the association between ΔDwell and choice consistency (Fig. 10F). The model of Callaway et al. (2021) predicts that ΔDwell depends on choice consistency, with greater ΔDwell for inconsistent choices. The explanation is the same as for the *aDDM*: if attention has a causal influence on choice, then for inconsistent choices it is highly likely that attention was directed to the item of lower value for a substantial fraction of the trial. Again, this is in clear contrast to what we observe in the data (Fig. 4).

We also performed simulations of the ‘optimal’ model of Jang et al. (2021), using the best-fitting parameters reported in their study (Fig. 10G). The model predicts that ΔDwell changes sign as a function of response time, contrary to what is observed in the data. For example, at response times of ∼1.5 s, it incorrectly predicts that people will tend to choose the option they looked at the least. The model of Jang et al. (2021) also predicts that the influence of the last fixation on choice depends on *Σr*, unlike what we observe in the data. We conclude that neither the *aDDM* nor the POMDP-based models account for the aspects of the data analyzed in Figs. 3 and 4.

## Discussion

We found post-decision signatures in the choice-RT-gaze data, suggesting that the association between gaze and choice, especially late in the trial, is not exclusively formative. Models positing an intra-decision (i.e., formative or constructive) multiplicative effect of attention on value—such as the attentional drift-diffusion model (*aDDM*)—predict that: (*i*) the last-fixation bias should be stronger when the items under consideration are overall more desirable, and (*ii*) ΔDwell—the difference in time spent looking at the chosen versus unchosen item—should be greater for inconsistent than for consistent choices. This is because inconsistent choices, in such models, benefit disproportionately from the attentional amplification of value. The data do not support these predictions (Fig. 3 and Fig. 4).

Instead, these observations are better explained by a *post-decision* account of the gaze-choice association—that is, one in which gaze shifts to the selected item *after* a covert commitment to a choice. We argue that directing gaze to the chosen item after a covert choice commitment is sensible, as the benefits of attending to a stimulus do not end with the decision itself. In naturalistic settings, for instance, selecting a food item is typically followed by the action of reaching toward it, where visual attention supports spatial localization and motor planning for the upcoming action. Although participants in our computerized task did not physically act on their choices, these sensorimotor processes are likely highly automatized and may still be engaged by default, even when not strictly required. Beyond motor preparation, post-decisional attention may also serve additional functions, such as facilitating sensory anticipation of the reward, supporting metacognitive evaluation of the decision, and contributing to value updating for future choices. From this perspective, a degree of attentional “stickiness”—whereby the chosen item remains preferentially attended after commitment—could emerge as an effectively optimal policy once these post-decisional processes are taken into account. Moreover, a specific feature of the task design may further reinforce this tendency: in the snacks paradigm, the unchosen item typically disappears from the screen immediately after a response is registered. It is therefore plausible that directing gaze to the chosen item after commitment partly reflects anticipation of the imminent disappearance of the unchosen option. To disentangle these mechanisms, it would be interesting for future work to test whether this attentional bias persists when the chosen item, rather than the unchosen one, is the stimulus that disappears upon response.

The *PDG* model captures many empirical observations, including the last-fixation bias and the gaze cascade effect. To account for the effect of overall value (*Σr*) on RT (Smith and Krajbich, 2019), the *PDG* model relies on the assumption that the variability of value representations increases with the values themselves (Ratcliff et al., 2018). This assumption is supported by neurobiological evidence. In the cortex, more desirable options tend to evoke higher firing rates (Platt and Glimcher, 1999; Padoa-Schioppa and Assad, 2006), and neural signal variance typically scales with its mean (Tolhurst et al., 1981). If the two value representations are independent, the variance of the momentary evidence, Δ*r*, should therefore increase in proportion to *Σr*. In contrast, the *aDDM* explains the same relation between overall value and RT more directly—as a consequence of the multiplicative effect of attention on value—and thus does so more parsimoniously (Krajbich et al., 2010). Moreover, empirical results show that higher overall value leads to both faster and more accurate choices, arguing against the notion that sensitivity decreases with value. Instead, participants appear to invest additional effort in high-value decisions, which may offset any increase in variability (Shevlin et al., 2022; Ting and Gluth, 2025).

A limitation of the *PDG* model is that underestimates the observed association between both early dwell-time and choice probability, as well as the association between ΔDwell and choice probability. Inspired by Westbrook et al. (2020), we therefore examined a *hybrid model* in which attention initially exerts a causal influence on the choice process and subsequently reflects the chosen option. This mixed model reintroduced some of the same limitations seen in the *aDDM*—most notably, its inability to account for the null effect of overall value on the last-fixation bias (MELFB) and the similarity in gaze bias across consistent and inconsistent decisions. We conclude that neither of these models is fully able to account for the data from the food-choice task.

The key assumption of the *PDG* model is that there is a delay between the moment a choice is internally committed and the moment it is externally reported with a key press. Because eye movements are typically faster than manual responses (*τ*_e_ < *τ*_m_ in our simulations), this delay creates a window during which gaze can already be directed toward the covertly chosen item before the response is formally registered. We do not interpret these non-decision latencies as irreducible physiological minima for moving the eyes or pressing a button (Bompas et al., 2025). Rather, they are inferred indirectly by fitting an additive non-decision-time parameter to the behavioral data, which we decompose into a sensory delay (*τ*_s_) and a manual execution delay (*τ*_m_). Values of *τ*_e_ are then chosen so that the model reproduces the observed magnitude of the behavioral effects. This estimation procedure has important limitations. Some participants show relatively “flat” chronometric functions: response times vary little with value despite otherwise normal psychometric performance. Such patterns likely reflect processes not explicitly represented in the model, including procrastination, reduced motivation, task-unrelated thought, or noise in item ratings. Within a drift-diffusion framework, however, these cases are accommodated by assigning a long non-decision time together with a short evidence-accumulation period (Table S1). Consequently, some estimated non-decision times are substantially longer than would be expected if they represented only sensory and motor delays. A further limitation is conceptual. We model non-decision time as occurring either before or after evidence accumulation, whereas in reality decisional and non-decisional components are likely temporally interleaved (Graziano et al., 2011). This simplification may also inflate the recovered latency estimates. With these caveats in mind, sensory and oculomotor delays on the order of 300 ms remain broadly plausible, although they likely lie near the upper end of a realistic range. The estimated eye-movement latency is especially long. For instance, in monkeys trained to report simple perceptual decisions with a saccade, roughly 100 ms elapses between the threshold-crossing signal in parietal cortex (or the superior colliculus) and the executed eye movement (Roitman and Shadlen, 2002; Stine et al., 2023). Crucially, however, varying the assumed non-decision latencies across a reasonable range does not alter the qualitative predictions of the model (Fig. 8).

Similar shortcomings to those observed in the *aDDM* were observed in models that derive attention-choice associations from optimal or near-optimal policies. Specifically, Callaway et al. (2021) and Jang et al. (2021) formalized the decision process as a partially observable Markov decision process (POMDP), in which attention enhances either the quantity (Callaway et al., 2021) or quality (Jang et al., 2021) of evidence about item value. To explain the gaze-choice association, both models assume that priors over item values are miscalibrated, such that decision-makers underestimate the true values. As a result, less-attended items are more biased toward zero, since less-attended items are more influenced by the miscalibrated prior. However, simulations based on these models fail to reproduce key empirical findings, including the association between ΔDwell, decision consistency, and RT (Fig. 10F,G).

The possibility that the gaze-choice association is partially post-decisional may help reconcile discrepancies between studies using free-viewing paradigms and those employing causal manipulations of attention. Common approaches to manipulating attention include limiting exposure duration (Frömer et al., 2022), interrupting trials based on gaze duration (Pärnamets et al., 2015; Newell and Le Pelley, 2018; Tavares et al., 2017; Pleskac et al., 2023), and cueing spatial attention (Störmer and Alvarez, 2016; Gwinn et al., 2019). A meta-analysis of the effects of visual attention on binary consumer choice (Bhatnagar and Orquin, 2022) found that these manipulations typically shift choice probabilities by ∼2–4% from a 50% baseline (cf., Tavares et al., 2017). These small effects contrast sharply with the large effects estimated by fits of the *aDDM*, which often posit a 70% discount of unattended items (Krajbich et al., 2010). The discrepancy between model predictions and the results of the attention-manipulation studies may arise from attempting to account for post-decision gaze-choice correlations using models that assume that the gaze-choice association is exclusively intra-decisional.

A post-decision gaze-choice association may offer a parsimonious explanation for differences in gaze behavior between tasks in which items are either chosen or rejected. In “choose” tasks, participants select the preferred item; in “reject” tasks, they exclude the less preferred item. Although logically equivalent in binary choice, the gaze is directed more to the preferred item in “choose” tasks and to the non-preferred item in “reject” tasks (Sepulveda et al., 2020; van der Laan et al., 2015; Mitsuda and Glaholt, 2014; Nittono and Wada, 2009). This difference has been explained by a variant of the *aDDM* in which attention modulates the integration of goal-relevant evidence (Sepulveda et al., 2020). Without ruling out this possibility, our results suggest that the difference in gaze allocation between “choose” and “reject” tasks arises because toward the end of the trial the gaze is directed to the selected option, regardless of whether it is to be accepted or rejected.

In this work, we focused on a simple yet widely used class of models in which decisions are based on comparing noisy value signals assigned to each item, and where explicit ratings are assumed to reflect the true underlying value of those items. An alternative class of models proposes that decisions arise from comparisons along individual feature dimensions, rather than at the level of the item as a whole (Tversky, 1972; Roe et al., 2001; Usher and McClelland, 2004; Busemeyer and Townsend, 1993; Summerfield and Tsetsos, 2015; Shadlen and Shohamy, 2016; Lichtenstein and Slovic, 2006; Johnson et al., 2007; Lee and Hare, 2023; Yang and Krajbich, 2023; Fisher, 2021). In food choice tasks, for example, relevant features might include expected satiety, caloric content, tastiness, saltiness, and so on (Rangel and Hare, 2010; Sullivan et al., 2015; Rramani et al., 2020; Suzuki et al., 2017). Attention is thought to fluctuate across these features, updating a decision variable for each alternative as different dimensions are sampled. Which dimensions are evaluated—and the weight assigned to each—may depend on the items being compared, past experience, or the broader decision context (Noguchi and Stewart, 2018; Roe et al., 2001; Juechems and Summerfield, 2019; Zylberberg et al., 2024; Lee and Pezzulo, 2022; Trueblood et al., 2014; Bhatia, 2013). As a result, the desirability of an item during the decision process may differ from that reported during the rating phase. A choice may appear inconsistent relative to initial ratings, but not relative to the specific features that were actually attended during deliberation. Similarly, if the ratings themselves are noisy and do not perfectly reflect subjective value, then some choices labeled as inconsistent may, in fact, be consistent with the decision-maker’s true preferences. Incorporating noise into the rating process would blur the distinction between consistent and inconsistent choices, potentially improving the alignment between model and data. These speculative ideas remain to be tested in future work.

Overall, our findings suggest that the association between attention and choice is not fully captured by models in which attention plays a purely causal role during decision formation. The data suggest an additional post-decision association, in which attention reflects, rather than shapes, the covert choice. While this does not preclude a formative influence of attention at earlier stages, it highlights the importance of considering the temporal dynamics of commitment when interpreting gaze patterns.

## Methods

### Food-choice task

We reanalyzed previously published data from six food-choice studies (Krajbich et al., 2010; Smith and Krajbich, 2018; Chen and Krajbich, 2016; Gwinn and Krajbich, 2016; Folke et al., 2016; Sepulveda et al., 2020). Across studies, the task followed the same two-phase paradigm. In the first (rating) phase, participants evaluated snack food items individually, either by rating how much they would like to consume each item on a numerical scale (Krajbich et al., 2010; Smith and Krajbich, 2018; Chen and Krajbich, 2016; Gwinn and Krajbich, 2016) or by stating their maximum willingness to pay using an incentive-compatible Becker–DeGroot–Marschak auction (Folke et al., 2016; Sepulveda et al., 2020). In the second (choice) phase, participants were presented with pairs of previously rated items and asked to choose which one they would prefer to consume at the end of the experiment, indicating their response with a key (or button) press. The studies differed in several design details. The number of items rated ranged from 70 (Krajbich et al., 2010) to 147 (Smith and Krajbich, 2018; Gwinn and Krajbich, 2016). Items with negative ratings were excluded from the choice phase in studies using a liking scale. The maximum permitted difference in value between the two items in a pair varied across studies: five rating points in Krajbich et al. (2010) and Smith and Krajbich (2018), three in Chen and Krajbich (2016), and one in Gwinn and Krajbich (2016); no such constraint was applied in Folke et al. (2016) and Sepulveda et al. (2020). The number of choice trials per participant ranged from 100 (Krajbich et al., 2010) to 240 (Sepulveda et al., 2020). In most studies, participants had to fixate on a central marker before each trial began. Further details of each experiment can be found in the original publications and in Table S5.

### *PDG* model

In the *PDG* model, choice and decision time are determined by the state of a scalar decision variable, *x*. The decision variable is the cumulative sum of samples from a normal distribution with mean *μdt* and variance σ^2^*dt*, and thus its evolution is described by the stochastic differential equation,

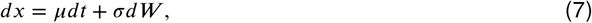

where *W* is the standard Wiener process and *x*(*t* = 0) = 0. The drift rate *μ* is given by

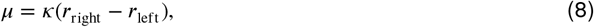

where *r*_left_ and *r*_right_ are the ratings assigned to the items presented on the left and right of the screen. The variance of the momentary evidence, σ^2^, scales linearly with the sum of the ratings (Eq. 5).

The accumulation process ends when the decision variable, *x*(*t*), reaches one of two bounds positioned symmetrically around zero, at ±*B*. The left (right) item is chosen when the decision variable reaches the upper (lower) bound. This first passage time establishes the decision time. The RT is the sum of the decision time plus the non-decision latencies, *τ*_nd_. We assume that *τ*_nd_ is Normally distributed with mean *μ*_nd_ and standard deviation σ_nd_.

The accumulation process starts after a short sensory delay, *τ*_s_ (Fig. 5), which is part of the non-decision latency. We used a fixed value of *τ*_s_ = 0.3 s for all participants. The remaining non-decision time (*τ*_m_ = *τ*_nd_ −*τ*_s_) is assigned to the time between crossing a decision bound and reporting the choice.

After crossing a bound, gaze is directed to the selected item. The time taken to switch gaze to the chosen item has an associated non-decision latency of *τ*_e_. This latency is assumed to follow a truncated Normal distribution with a mean of *μ*_*e*_ = 0.35 seconds and a standard deviation σ_*e*_ of one-third of the mean, truncated to ensure non-negative values. Since *τ*_e_ is usually smaller than *τ*_m_, the gaze is informative about the choice.

We fit the model parameters λ = {*κ, B, γ, μ*_nd_, σ_nd_} using maximum likelihood to the choice and RT data from each trial. Fits were performed independently for each participant. The log-likelihood of the parameters is given by:

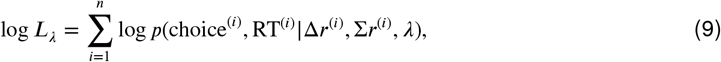

where the summation is over trials. The joint probability of choice and decision time was obtained by numerically solving the Fokker-Planck equation associated with the drift-diffusion process using the Chang-Cooper fully implicit method (Chang and Cooper, 1970; Zylberberg et al., 2016; Kiani and Shadlen, 2009). To obtain the joint probability distribution over choice and RT, we convolve the probability distribution of decision times with the distribution of non-decision times, given by a truncated normal distribution with parameters *μ*_nd_ and σ_nd_ (the truncation constrains the non-decision times to be positive).

## Attentional drift-diffusion model

We simulated the *aDDM* (Eqs. 1 and 2) using the best-fitting parameters reported by Krajbich et al. (2010): *d* = 0.0002 ms^−1^, σ = 0.02, *θ* = 0.3 and bounds *B* = ±1. The response time was calculated as the sum of the decision time from the drift-diffusion process and a fixed non-decision time of *t*_nd_ = 0.355 s, as in Smith and Krajbich (2019).

In the *aDDM* (as well as in the *DDM*), it is necessary to set one of the parameters *d*, σ, or *B* to a fixed value for the model parameters to be identifiable. In the original description of the *aDDM* (Krajbich et al., 2010), the upper and lower bounds were set to a fixed value of ±1. For consistency with the *PDG* model, we reformulated the *aDDM* so that the variance of the noise accumulated during one second of unbounded accumulation is equal to 1. Then the decision variable of the *aDDM, x*(*t*), evolves as in Eq. 7 with σ = 1. Time is discretized in steps of dt = 0.001 s. The drift rate *μ* is equal to *κ*(*r*_right_ − *θr*_left_) when looking at the item on the right, and to *κ*(*θr*_right_ − *r*_left_) when looking at the item on the left, where *κ* is the signal-to-noise. A model equivalent to the original *aDDM* is obtained with the following parameters:

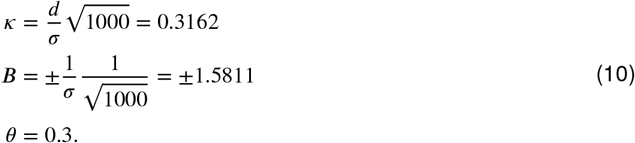

Simulating the *aDDM* requires modeling how attention alternates between the two items. We fit the empirically-observed dwell durations with log-normal distributions. Separate fits were conducted for the first and middle dwells. Middle dwells are those that were neither the first nor the last of the stimulus-viewing epoch. The fits provide a good match to the experimental data (Fig. S2). Each simulated trial of the *aDDM* begins by sampling from the first dwell distribution, followed by sampling from the middle dwell distribution until a decision threshold is reached. Similar to the experimental data, the first fixation has a 0.74 probability of being directed to the left item, and attention alternates between the two items thereafter. We simulated the same trials (same value pairs) as those completed by the participants, repeating each trial 10 times.

We fit three variants of the *aDDM* to individual participant data. In the first variant, attention has an additive effect on value, rather than a multiplicative effect (Fig. 10B); see equations for the drift rate in the main text). In the second variant (Fig. 10D), the drift rate exhibits inter-trial variability. Specifically, on each trial, the drift rate *μ* is given by *κ*(*r*_left_ −*θr*_right_) +*v* when fixating the left item, and by *κ*(*θr*_left_ −*r*_right_) +*v* when fixating the right item. Here, *v* is trial-specific noise drawn from a normal distribution with mean 0 and standard deviation σ_*d*_, which is estimated from the data. Importantly, *v* is constant within a trial but varies across trials.

The third *aDDM* variant can be interpreted as alternative implementation of the one with inter-trial drift-rate variability. The noise does not directly affect the drift rate, but the items’ value. Specifically, the drift rate *μ* is given by *κ*((*r*_left_ + *v*_1_) − *θ*(*r*_right_ + *v*_2_)) when fixating the left item, and by *κ*(*θ*(*r*_left_ + *v*_1_) − (*r*_right_ + *v*_2_)) when fixating the right item. Here, *v*_*x*_ represents trial-specific noise, drawn from a normal distribution with mean 0 and standard deviation σ_*d*_ . This formulation reflects the idea that the value reported in the rating phase may deviate from the item’s “true” underlying value (c.f., Zylberberg et al. (2024); Polania et al. (2019)).

Additionally, we simulated the *aDDM* using the best-fitting parameters reported in previous studies by Krajbich et al. (2010) and Smith and Krajbich (2019) (Fig. S3).

The core parameters of the *aDDM* are λ = {*κ, B, θ*}, corresponding respectively to the signal-to-noise ratio, the decision bound height, and the value-scaling factor applied to the unattended option. In the model with inter-trial drift-rate variability, σ_*d*_ is added to capture the standard deviation of the drift-rate noise. The models are fit to maximize the likelihood of the parameters given the choice and decision time (DT), and given the sequence and duration of the dwells observed on each trial:

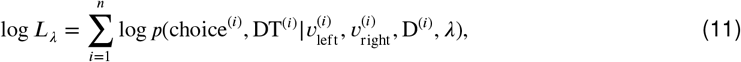

where D^(*i*)^ is the sequence and duration of dwells observed on trial *i*, and DT^(*i*)^ is the decision time on trial *i* that is assumed to be equal to the sum of the dwells on either of the two items. The joint probability of choice and decision time was computed via numerical approximations of the corresponding Fokker-Planck (FP) equation. Unlike the special case where *θ* = 1, here the drift rate varies with the focus of attention. As a result, solving the FP equation becomes more computationally demanding, since the probability density of the decision variable must be computed separately for each trial, given the trial-specific stochastic fluctuations in gaze. Numerical solutions were obtained using a fully implicit method (Chang and Cooper, 1970), propagating the probability density of the decision variable over time adjusting the drift rate depending on the focus of gaze.

For the model with inter-trial variability in the drift-rate, we discretized the distribution of drift perturbations into a finite number of bins. Specifically, we drew *n*_bins_=11 quantile-based samples from a zero-mean Gaussian distribution with standard deviation σ_drift_, using the midpoint of each quantile as a representative value. We numerically solved the Fokker–Planck equation independently for each of these bins. The final probability distribution over choice and decision times was computed as the average (uniform-weighted) across the solutions for each bin, effectively marginalizing over the distribution of drift perturbations.

We use the best-fitting parameters to simulate the *aDDM* independently for each participant. From the simulations we obtain a choice and decision time for each trial. Response times are estimated as the decision time plus mean non decision time, *μ*_*nd*_, defined as the trial-average RT of each participant minus the trial-average decision time obtained from the model simulations.

### Model simulations

Simulations of the *PDG* model (Fig. 6) and *aDDM* (Fig. 10A–E) were made using the same trials (same value pairs) that participants completed, with each trial repeated 10 times. Dwell durations were randomly sampled from log-Normal distributions fit to the duration of the dwells (Fig. S2). First and subsequent dwells were fit separately. The probability of sampling the left item first was set to 0.74 to match the value obtained from the experimental data.

### Combined *aDDM***-***PDG* model

The combined *aDDM*-*PDG* model (Fig. 10C) builds on the *aDDM* fit to single-participant data. We add to the data simulated with the *aDDM* (Fig. 10A) two aspects of the *PDG* model: (i) a sensory delay between the onset of the food items and the start of the evidence accumulation process of *τ*_s_, and (ii) another delay *τ*_e_ between the time at which a bound is crossed and the time at which the gaze is directed to the chosen item (truncated such that *τ*_e_ is non-negative). Parameters *τ*_s_, *μ*_*e*_ and σ_*e*_ were set to 0.25s, 0.2s and 0.05s respectively. The same approach and parameter values were used in the *aDDM* model with inter-trial variability in the drift rate (Fig. 10E).

### Model fitting

Parameter optimization was performed using the Bayesian Adaptive Direct Search (BADS) method (Acerbi and Ma, 2017). Table S1, Table S2, Table S3 and Table S4 show the best-fitting parameters for the *PDG* model, the *aDDM*, the model with intra-decisional additive attention, and the *aDDM* with inter-trial drift-rate variability, respectively.

### Optimal decision models

The optimal model of Jang et al. (2021) has 5 free parameters: the cost of switching attention between items (*c*_*s*_), the cost, per second, of accumulating evidence (*c*), the variance of the evidence sampling distribution 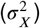, the variance of the prior distribution 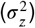, and the relative information gain for attended vs. unattended items (*κ*). See Jang et al. (2021) for a detailed explanation of the model. We simulated 1280 trials per participant with the parameters that Jang et al. (2021) reported best replicated the human behavioral data: *c*_*s*_ = 0.0065, *c* = 0.23, 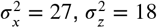, *κ* = 0.004. As in Jang et al., the prior mean over the items’ values, 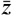, was set to zero, which is lower than the true mean value of the items. The model requires this feature to produce a gaze bias (Jang et al., 2021).

We also analyzed the simulations of the optimal model developed by Callaway et al. (2021). The model has five free parameters: the standard deviation of the evidence sampling distribution, the cost of obtaining a sample, the cost of switching attention between items, the degree to which the prior over the item values is biased toward zero, and a ‘temperature’ parameter of a Boltzmann distribution, which controls the degree of stochasticity in the selection of the optimal policy. The simulations (N = 4,550,400 trials) with the parameters that best fit the human behavioral data were kindly provided by Frederick Callaway and are available at https://github.com/fredcallaway/optimal-fixations-simple-choice. A detailed explanation of the model and the fitting procedure can be found in the original publication (Callaway et al., 2021).

## Data analysis

The dashed lines in Fig. 3(top) were derived from the following logistic regression model:

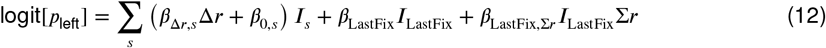

*I*_LastFix_ takes a value of 1 if the left item was fixated on last and 0 if the right item was fixated last. The rightmost term captures the interaction of *I*_LastFix_ with *Σr*; the associated *β* is the slope of the dashed line in Fig. 3(top).

We fit the following linear regression model to test for an association between choice consistency (*c*) and the difference in looking time between the chosen and unchosen items (ΔDwell):

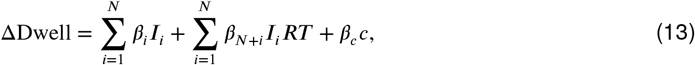

where *N* is the number of participants, *I*_*i*_ is an indicator variable that takes the value 1 if the trial was completed by subject *i* and 0 otherwise, and *c* is equal to 1 for trials in which the higher-rated item was chosen, defined as a *consistent* choice, and 0 for trials in which the lower-rated item was chosen, defined as an *inconsistent* choice. We used a one-tailed *t* -test to test whether *β*_*c*_ is negative, that is, whether ΔDwell is larger for inconsistent than for consistent choices. When the use of a one-tailed test is not explicitly mentioned, we used two-tailed *t* -tests to test whether *β*_*c*_ differed significantly from zero.

To test the hypothesis that there was no difference in the probability of looking at the chosen item before and after the choice report (Fig. 7), we used a Wilcoxon signed-rank test. We define two time epochs, one from -200 ms to 0 ms, and the other from 0 ms to 200 ms, relative to the time of the choice report. For each time epoch, we calculated the time each participant spent looking at the chosen item and divided it by the time spent looking at either item. We obtain two proportions per participant (Fig. 7B), which we subjected to a two-tailed Wilcoxon signed-rank test.

For the plots showing the probability of looking at the chosen item aligned to stimulus onset (e.g., Fig. 6E, left), we eliminated the gaze information from the 500 ms prior to the choice report. This step was taken to eliminate, from the stimulus-aligned plots, gaze effects that might be related to the response.

For the plots showing the psychometric function split by whether the last fixated item was the one on the left or right (e.g., Fig. 6D), we classified trials as ‘left item fixated on last’ and ‘right item fixated on last’ depending on which item was being looked at at the time of the choice report. We excluded trials in which the participant either was not fixating on one of the two relevant items at the time of the choice report or the direction of gaze could not be resolved (e.g., eye blinks). We repeated the analyses reclassifying trials according to which item was last looked at before the choice report regardless of when it occurred during the trial, and obtained nearly identical results.

To test the association between first-dwell duration and the probability of choosing the option that was fixated first (Fig. 9), we fit the following logistic regression model, separately for each total–dwell–count condition (2–5 total dwells):

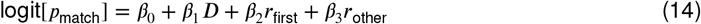

where *p*_match_ is the probability of choosing the item fixated first, *D* is the standardized first-dwell duration, and *r*_first_ and *r*_other_ represent the value ratings of the first-fixated and alternative items, respectively. Statistical significance of the first-dwell-duration coefficient was assessed using a likelihood-ratio test, comparing the full model to a reduced model that excluded the *β*_1_ term.

To illustrate the gaze cascade effect (e.g., Fig. 6E), we filled a matrix of dimensions `NumberOfTrials × NumberOfTimeSteps,` with a 1 when participants were looking at the right item, a 0 when they were looking at the left item, and a NaN otherwise. The time step was 1 millisecond. We averaged this matrix across trials, ignoring the NaNs, first within participants and then across participants. For the RT aligned plots, we followed the same procedure after aligning each trial to the response time. For the *aDDM* and Callaway’s optimal model, we followed a slightly different procedure, because in these models the drift-rate is undefined when participants are not looking at either of the two items. Therefore, we removed from each trial the times when the participant’s gaze was not directed to either snack item, and aligned the responses to the total fixation time rather than to the response time.

## Data Availability

Code and data required to reproduce the model fitting, simulations, and figures presented in this paper are available at https://github.com/arielzylberberg/PostDecisionalAttention_eLife2026.

## Acknowledgments

We are grateful to Fred Callaway and Yaniv Abir for helpful comments on an earlier version of the manuscript, to Fred Callaway for kindly sharing the simulations of his model, and to Antonio Rangel for sharing data and for helpful and stimulating discussions.

## Supplemental information

**Figure S1.**
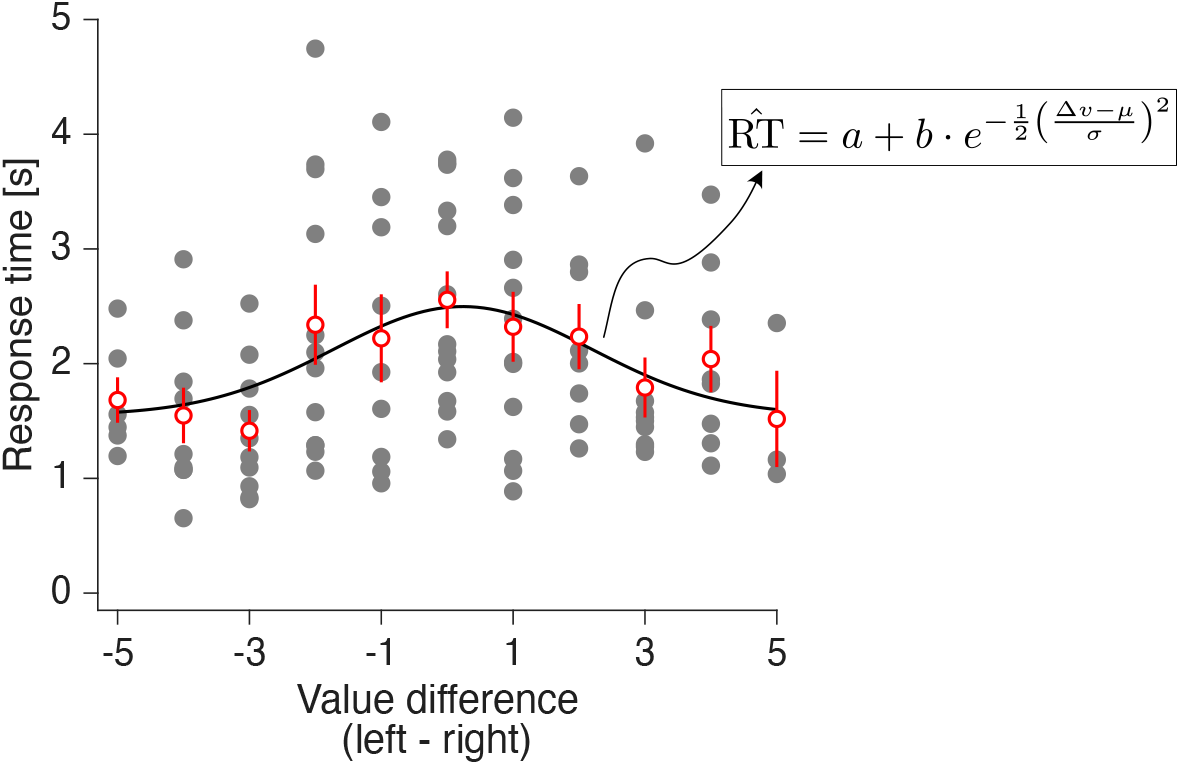
Removing the contribution of Δ*r* from response times. Illustration of the method used to remove the contribution of Δ*r* from the RT. The gray markers indicate the RT for each trial of a representative participant. The abscissa are the values of Δ*r* for the corresponding trial. These data points were fitted with the bell-shaped function shown in the figure, with parameters *a, b, μ* and σ. The best-fitting function captures the general trend in the data, as can be seen by comparing the model fits (black solid line) with the average response time per value of Δ*r* (red, mean plus s.e.m.). To compute the RT residuals, we subtract from each trial the value of 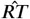 corresponding to the corresponding value of Δ*r*. Fits were performed independently for each participant.

**Figure S2.**
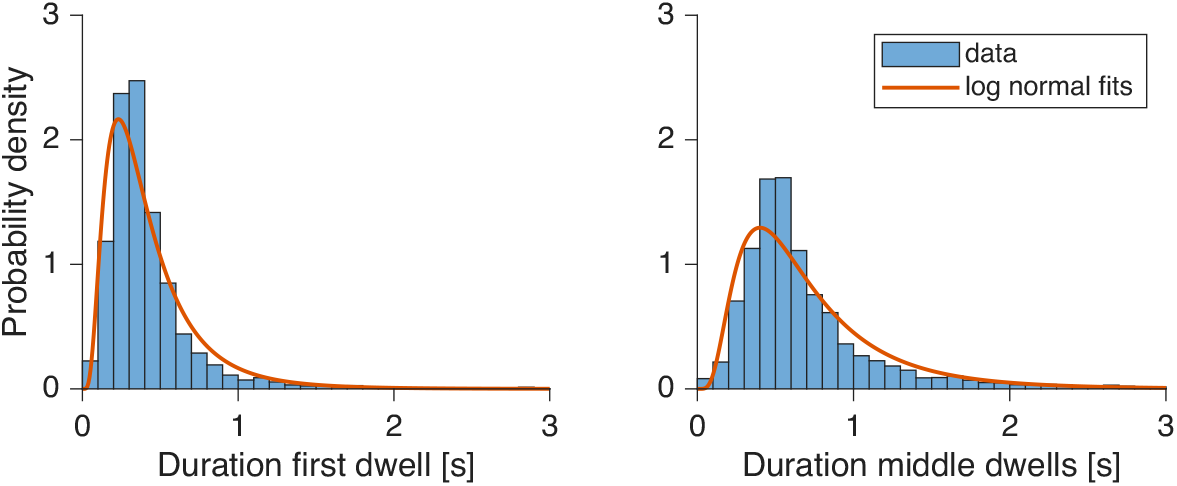
Fits of the duration of the dwells. Distribution of the durations of the first dwell (left) and middle dwells (right). Middle dwells include all dwells except the first and last. The durations were fitted with a log-normal distribution (red), independently for the first and middle dwells. The best-fitting log-normal parameters were used to simulate the *aDDM* and *PDG*. On each trial, the first dwell is sampled from the corresponding distribution, and the subsequent dwell durations up to the bound crossing are sampled from the distribution of middle dwells.

**Figure S3.**
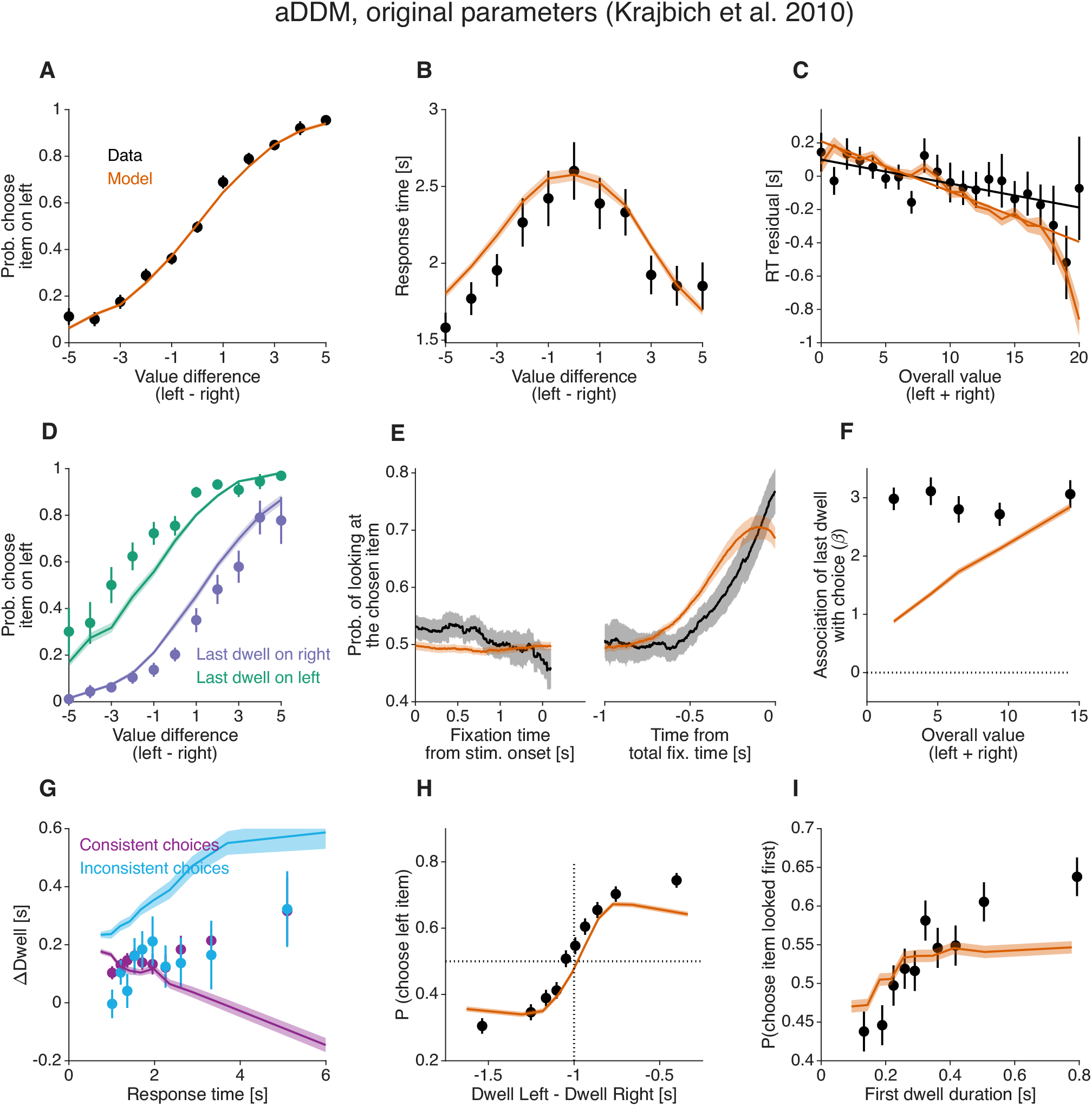
*aDDM* model. Model simulations were generated using the *aDDM* with the best-fitting parameters reported by Krajbich et al. (2010) and Smith and Krajbich (2019). This version of the model assumes constant (i.e., flat) decision bounds and no inter-trial variability in drift rate. Unlike the models presented in the main text, it was fit by Krajbich et al. (2010) to data pooled across participants. Same conventions as in Fig. 6.

**Figure S4.**
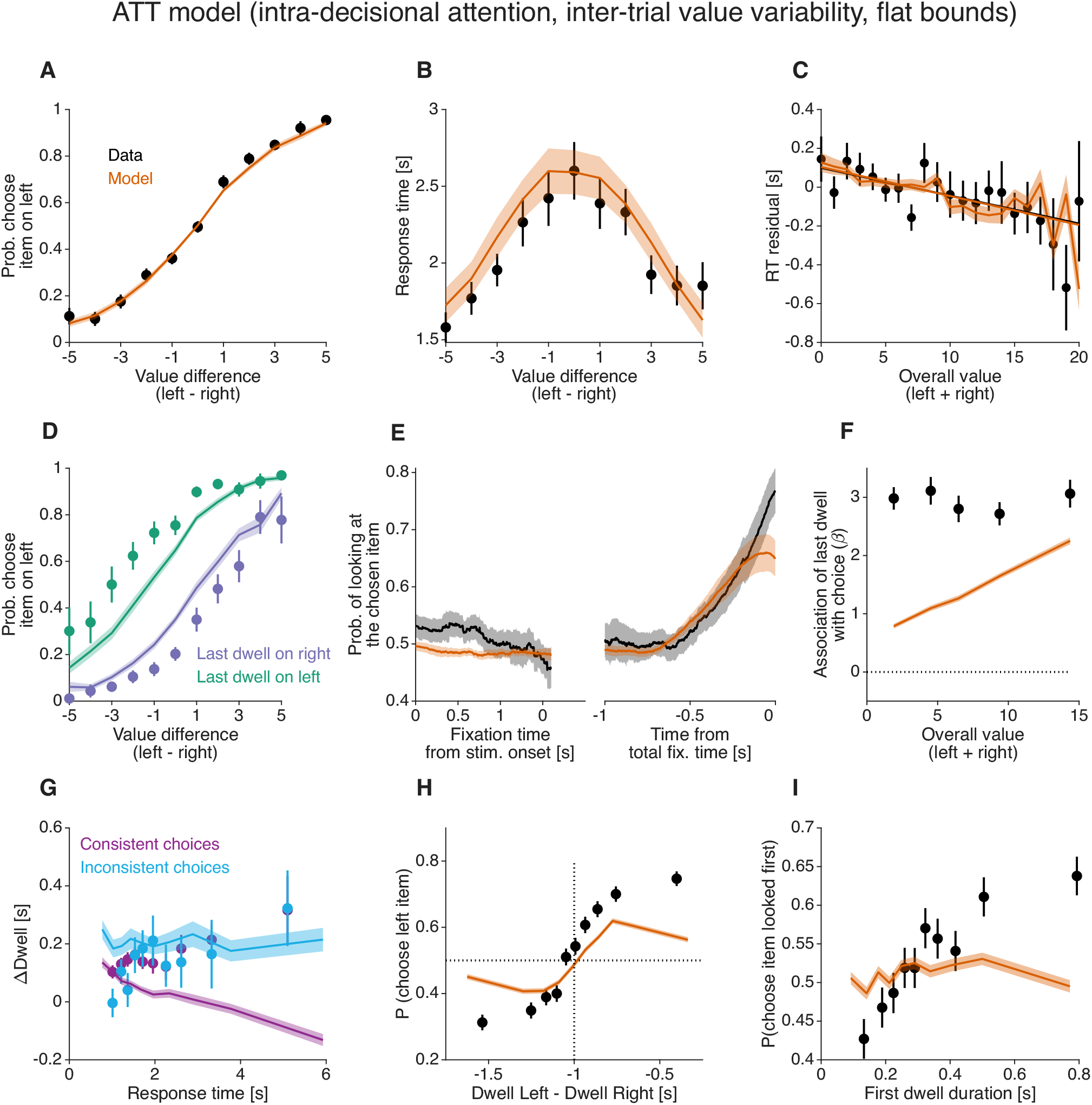
Model with intra- and post-decisional attention, and inter-trial variability in the items’ value. Unlike the model illustrated in Fig. 10D, here the items’ values—instead of the drift rates—are corrupted with additive Gaussian noise. This noise is fixed within each trial but varies randomly across trials. Model parameters were fit to individual participants’ choice, RT, and fixation data. Same conventions as in Fig. 6.

**Table S1.** Best-fitting parameter values for the *PDG* model.

| Subject | $\kappa$ | $\mu_{nd}$ | $\sigma_{nd}$ | $B$ | $\gamma$ |
| --- | --- | --- | --- | --- | --- |
| 1 | 0.369 | 0.536 | 0.018 | 1.293 | 0.000 |
| 2 | 0.076 | 0.671 | 0.091 | 1.311 | 0.067 |
| 3 | 0.187 | 0.600 | 0.144 | 2.084 | 0.000 |
| 4 | 0.216 | 1.219 | 0.235 | 0.999 | 0.141 |
| 5 | 0.383 | 0.544 | 0.010 | 2.004 | 0.171 |
| 6 | 0.244 | 0.886 | 0.140 | 0.613 | 0.000 |
| 7 | 0.197 | 0.536 | 0.035 | 2.116 | 0.055 |
| 8 | 0.617 | 1.090 | 0.143 | 1.612 | 0.247 |
| 9 | 0.551 | 0.702 | 0.044 | 1.254 | 0.137 |
| 10 | 0.292 | 0.706 | 0.142 | 1.483 | 0.038 |
| 11 | 0.353 | 0.730 | 0.012 | 1.467 | 0.000 |
| 12 | 0.243 | 0.525 | 0.010 | 1.547 | 0.000 |
| 13 | 0.242 | 0.646 | 0.013 | 1.707 | 0.021 |
| 14 | 0.332 | 0.649 | 0.011 | 1.201 | 0.000 |
| 15 | 0.058 | 0.598 | 0.010 | 2.184 | 0.233 |
| 16 | 0.335 | 0.738 | 0.143 | 1.065 | 0.000 |
| 17 | 0.224 | 0.774 | 0.010 | 2.150 | 0.024 |
| 18 | 0.198 | 1.498 | 0.338 | 1.810 | 0.021 |
| 19 | 0.459 | 0.551 | 0.010 | 2.487 | 0.212 |
| 20 | 0.192 | 0.778 | 0.163 | 1.673 | 0.045 |
| 21 | 0.349 | 0.523 | 0.010 | 2.079 | 0.296 |
| 22 | 0.425 | 1.255 | 0.195 | 1.298 | 0.009 |
| 23 | 0.535 | 0.879 | 0.108 | 0.994 | 0.180 |
| 24 | 0.314 | 1.225 | 0.222 | 1.024 | 0.061 |
| 25 | 0.348 | 0.973 | 0.129 | 0.888 | 0.025 |
| 26 | 0.203 | 0.626 | 0.017 | 1.243 | 0.000 |
| 27 | 0.468 | 0.538 | 0.047 | 0.862 | 0.145 |
| 28 | 0.057 | 1.071 | 0.010 | 1.150 | 0.002 |
| 29 | 0.258 | 0.654 | 0.023 | 1.916 | 0.047 |
| 30 | 0.291 | 0.680 | 0.182 | 1.448 | 0.000 |
| 31 | 0.363 | 0.540 | 0.010 | 1.438 | 0.176 |
| 32 | 0.340 | 0.679 | 0.017 | 1.622 | 0.076 |
| 33 | 0.450 | 0.880 | 0.163 | 0.702 | 0.000 |
| 34 | 0.439 | 0.944 | 0.145 | 1.110 | 0.000 |
| 35 | 0.418 | 0.615 | 0.086 | 1.116 | 0.320 |
| 36 | 0.281 | 0.735 | 0.178 | 1.057 | 0.095 |
| 37 | 0.066 | 0.859 | 0.156 | 0.866 | 0.000 |
| 38 | 0.268 | 0.722 | 0.014 | 1.668 | 0.004 |
| 39 | 0.259 | 0.662 | 0.034 | 1.336 | 0.007 |

**Table S2.**
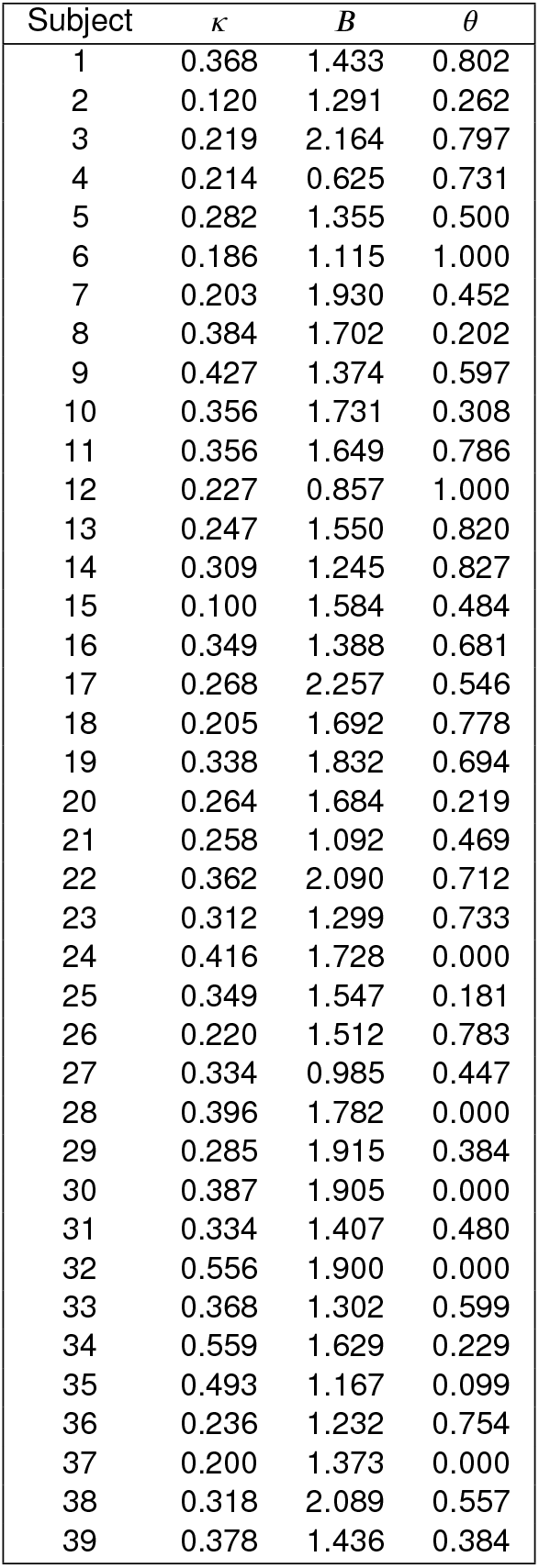
Best-fitting parameter values for the *aDDM*.

**Table S3.** Best-fitting parameter values for the model with additive intra-decision attention.

| Subject | $\kappa$ | $B$ | $\omega$ |
| --- | --- | --- | --- |
| 1 | 0.334 | 1.442 | 1.511 |
| 2 | 0.100 | 1.294 | 5.000 |
| 3 | 0.197 | 2.171 | 0.700 |
| 4 | 0.185 | 0.625 | 0.866 |
| 5 | 0.218 | 1.338 | 3.531 |
| 6 | 0.186 | 1.115 | -0.820 |
| 7 | 0.152 | 1.927 | 3.584 |
| 8 | 0.285 | 1.730 | 5.000 |
| 9 | 0.349 | 1.399 | 2.522 |
| 10 | 0.261 | 1.731 | 5.000 |
| 11 | 0.320 | 1.658 | 1.499 |
| 12 | 0.238 | 0.858 | -0.620 |
| 13 | 0.225 | 1.545 | 0.302 |
| 14 | 0.281 | 1.245 | 0.869 |
| 15 | 0.100 | 1.586 | 0.864 |
| 16 | 0.293 | 1.430 | 2.897 |
| 17 | 0.213 | 2.279 | 3.123 |
| 18 | 0.183 | 1.685 | 1.289 |
| 19 | 0.288 | 1.830 | 0.957 |
| 20 | 0.163 | 1.684 | 5.000 |
| 21 | 0.195 | 1.090 | 4.289 |
| 22 | 0.307 | 2.099 | 2.395 |
| 23 | 0.272 | 1.346 | 2.682 |
| 24 | 0.249 | 1.752 | 5.000 |
| 25 | 0.233 | 1.577 | 5.000 |
| 26 | 0.196 | 1.515 | 1.241 |
| 27 | 0.238 | 0.980 | 2.740 |
| 28 | 0.209 | 1.716 | 5.000 |
| 29 | 0.202 | 1.930 | 3.166 |
| 30 | 0.272 | 1.950 | 5.000 |
| 31 | 0.255 | 1.457 | 2.967 |
| 32 | 0.288 | 1.910 | 3.207 |
| 33 | 0.283 | 1.329 | 3.061 |
| 34 | 0.353 | 1.635 | 2.081 |
| 35 | 0.254 | 1.162 | 5.000 |
| 36 | 0.207 | 1.236 | 1.538 |
| 37 | 0.113 | 1.308 | 5.000 |
| 38 | 0.246 | 2.105 | 3.384 |
| 39 | 0.257 | 1.443 | 1.841 |

**Table S4.** Best-fitting parameter values for the *aDDM* with inter-trial drift-rate variability.

| Subject | $\kappa$ | $B$ | $\theta$ | $\sigma_{\text{drift}}$ |
| --- | --- | --- | --- | --- |
| 1 | 0.369 | 1.435 | 0.803 | 0.000 |
| 2 | 0.158 | 1.495 | 0.405 | 0.804 |
| 3 | 0.223 | 2.179 | 0.799 | 0.095 |
| 4 | 0.213 | 0.625 | 0.727 | 0.000 |
| 5 | 0.464 | 1.804 | 0.576 | 1.205 |
| 6 | 0.335 | 1.762 | 1.000 | 1.494 |
| 7 | 0.263 | 2.172 | 0.532 | 0.490 |
| 8 | 0.570 | 2.514 | 0.224 | 1.125 |
| 9 | 0.430 | 1.377 | 0.596 | 0.091 |
| 10 | 0.578 | 2.463 | 0.415 | 1.062 |
| 11 | 0.494 | 2.041 | 0.791 | 0.808 |
| 12 | 0.227 | 0.855 | 1.000 | 0.007 |
| 13 | 0.495 | 2.075 | 0.877 | 1.050 |
| 14 | 0.387 | 1.356 | 0.868 | 0.621 |
| 15 | 0.100 | 1.788 | 0.634 | 0.681 |
| 16 | 0.373 | 1.439 | 0.678 | 0.336 |
| 17 | 0.871 | 5.911 | 0.590 | 2.128 |
| 18 | 0.204 | 1.690 | 0.778 | 0.000 |
| 19 | 0.861 | 3.409 | 0.669 | 1.543 |
| 20 | 0.385 | 2.144 | 0.293 | 0.816 |
| 21 | 0.258 | 1.090 | 0.468 | 0.007 |
| 22 | 1.160 | 6.722 | 0.721 | 2.582 |
| 23 | 0.381 | 1.553 | 0.772 | 0.822 |
| 24 | 0.854 | 3.662 | 0.000 | 1.740 |
| 25 | 0.466 | 2.039 | 0.235 | 0.927 |
| 26 | 0.422 | 2.157 | 0.803 | 1.178 |
| 27 | 0.378 | 1.101 | 0.477 | 0.815 |
| 28 | 0.686 | 2.889 | 0.000 | 1.361 |
| 29 | 0.357 | 2.164 | 0.443 | 0.522 |
| 30 | 0.551 | 2.482 | 0.000 | 0.886 |
| 31 | 0.502 | 1.822 | 0.563 | 1.000 |
| 32 | 0.839 | 2.622 | 0.114 | 0.884 |
| 33 | 0.512 | 1.763 | 0.641 | 1.057 |
| 34 | 0.708 | 1.968 | 0.301 | 0.688 |
| 35 | 0.856 | 1.827 | 0.260 | 1.555 |
| 36 | 0.309 | 1.410 | 0.778 | 0.738 |
| 37 | 0.384 | 2.248 | 0.000 | 1.460 |
| 38 | 0.445 | 2.581 | 0.604 | 0.637 |
| 39 | 0.628 | 1.887 | 0.447 | 1.062 |

**Table S5.** Comparison of experimental design features across food choice studies.

| Feature | Krajbich et al.<br>(2010) | Smith &<br>Krajbich (2018) | Chen &<br>Krajbich (2016) | Gwinn &<br>Krajbich (2016) | Folke et al.<br>(2016) | Sepulveda et al.<br>(2020) |
| --- | --- | --- | --- | --- | --- | --- |
| Number of subjects | 39 | 44 | 44 | 36 | 28 | 31 |
| Valuation method | Liking (−10/+10) | Liking (−10/+10) | Liking (−10/+10) | Yes/No + Liking (0–10) | BDM (£0–£3) | BDM (£0–£3) |
| Items rated | 70 | 147 | 139 | 147 | 60 | 60 |
| Max value difference | 5 | 5 | 3 | 1 | Unconstrained | Unconstrained |
| Number of trials | 100 | 200 | 200 | 200 | 120 | 240 |
| Confidence rating | No | No | No | No | Yes | Yes |
| Task framing | Like only | Like only | Like only | Like only | Like only | Like + Dislike |
| Post-session re-rating | No | No | No | Yes | No | No |

## References

Acerbi L, Ma WJ. Practical Bayesian optimization for model fitting with Bayesian adaptive direct search. arXiv preprint arXiv:170504405. 2017; .

Armel KC, Beaumel A, Rangel A. Biasing simple choices by manipulating relative visual attention. Judgment and Decision making. 2008; 3(5):396–403.

Bhatia S. Associations and the accumulation of preference. Psychological review. 2013; 120(3):522.

Bhatnagar R, Orquin JL. A meta-analysis on the effect of visual attention on choice. Journal of Experimental Psychology: General. 2022; .

Bompas A, Sumner P, Hedge C. Non-decision time: The Higgs Boson of decision. Psychological Review. 2025; 132(2):330.

Busemeyer JR, Townsend JT. Decision field theory: a dynamic-cognitive approach to decision making in an uncertain environment. Psychological review. 1993; 100(3):432.

Callaway F, Rangel A, Griffiths TL. Fixation patterns in simple choice reflect optimal information sampling. PLoS computational biology. 2021; 17(3):e1008863.

Cavanagh JF, Wiecki TV, Kochar A, Frank MJ. Eye tracking and pupillometry are indicators of dissociable latent decision processes. Journal of Experimental Psychology: General. 2014; 143(4):1476.

Chang J, Cooper G. A practical difference scheme for Fokker-Planck equations. Journal of Computational Physics. 1970; 6(1):1–16.

Chen WJ, Krajbich I. Pupil dilation and attention in value-based choice. Unpublished manuscript, The Ohio State University. 2016; .

Fisher G. A multiattribute attentional drift diffusion model. Organizational Behavior and Human Decision Processes. b2021; 165:167–182.

Folke T, Jacobsen C, Fleming SM, De Martino B. Explicit representation of confidence informs future value-based decisions. Nature Human Behaviour. 2016; 1(1):0002.

Frömer R, Callaway F, Griffiths T, Shenhav A. Considering what we know and what we don’t know: Expectations and confidence guide value integration in value-based decision-making. PsyArXiv. 2022; .

Glaholt MG, Reingold EM. The time course of gaze bias in visual decision tasks. Visual Cognition. 2009; 17(8):1228– 1243.

Gluth S, Deakin J, Rieskamp J. A theory of multiattribute search and choice. Psychological review. 2026; .

Gluth S, Kern N, Kortmann M, Vitali CL. Value-based attention but not divisive normalization influences decisions with multiple alternatives. Nature human behaviour. 2020; 4(6):634–645.

Graziano M, Polosecki P, Shalom DE, Sigman M. Parsing a perceptual decision into a sequence of moments of thought. Frontiers in integrative neuroscience. 2011; 5:45.

Gwinn R, Leber AB, Krajbich I. The spillover effects of attentional learning on value-based choice. Cognition. 2019; 182:294–306.

Gwinn RE, Krajbich I, Attitudes and attention: How attitude accessibility and certainty influence attention and subjective choice; 2016.

Hébert B, Woodford M. Information costs and sequential information sampling. National Bureau of Economic Research; 2018.

Jang AI, Sharma R, Drugowitsch J. Optimal policy for attention-modulated decisions explains human fixation behavior. Elife. 2021; 10:e63436.

Johnson EJ, Häubl G, Keinan A. Aspects of endowment: a query theory of value construction. Journal of experimental psychology: Learning, memory, and cognition. 2007; 33(3):461.

Juechems K, Summerfield C. Where does value come from? Trends in cognitive sciences. 2019; 23(10):836–850.

Kiani R, Shadlen MN. Representation of confidence associated with a decision by neurons in the parietal cortex. science. 2009; 324(5928):759–764.

Krajbich I. Accounting for attention in sequential sampling models of decision making. Current opinion in psychology. 2019; 29:6–11.

Krajbich I, Armel C, Rangel A. Visual fixations and the computation and comparison of value in simple choice. Nature neuroscience. 2010; 13(10):1292–1298.

Krajbich I, Lu D, Camerer C, Rangel A. The attentional drift-diffusion model extends to simple purchasing decisions. Frontiers in psychology. 2012; 3:193.

Krajbich I, Rangel A. Multialternative drift-diffusion model predicts the relationship between visual fixations and choice in value-based decisions. Proceedings of the National Academy of Sciences. 2011; 108(33):13852–13857.

van der Laan LN, Hooge IT, De Ridder DT, Viergever MA, Smeets PA. Do you like what you see? The role of first fixation and total fixation duration in consumer choice. Food Quality and Preference. 2015; 39:46–55.

Lee DG, Hare TA. Evidence accumulates for individual attributes during value-based decisions. Decision. 2023; 10(4):330.

Lee DG, Pezzulo G. Choice-induced preference change under a sequential sampling model framework. bioRxiv. 2022; p. 2022–07.

Lichtenstein S, Slovic P. The construction of preference. Cambridge University Press; 2006.

Link SW. The relative judgment theory of two choice response time. Journal of Mathematical Psychology. 1975; 12(1):114–135.

Mitsuda T, Glaholt MG. Gaze bias during visual preference judgements: Effects of stimulus category and decision instructions. Visual Cognition. 2014; 22(1):11–29.

Newell BR, Le Pelley ME. Perceptual but not complex moral judgments can be biased by exploiting the dynamics of eye-gaze. Journal of Experimental Psychology: General. 2018; 147(3):409.

Nittono H, Wada Y. Gaze shifts do not affect preference judgments of graphic patterns. Perceptual and motor skills. 2009; 109(1):79–94.

Noguchi T, Stewart N. Multialternative decision by sampling: A model of decision making constrained by process data. Psychological review. 2018; 125(4):512.

Padoa-Schioppa C, Assad JA. Neurons in the orbitofrontal cortex encode economic value. Nature. 2006; 441(7090):223–226.

Pärnamets P, Johansson P, Hall L, Balkenius C, Spivey MJ, Richardson DC. Biasing moral decisions by exploiting the dynamics of eye gaze. Proceedings of the National Academy of Sciences. 2015; 112(13):4170–4175.

Platt ML, Glimcher PW. Neural correlates of decision variables in parietal cortex. Nature. 1999; 400(6741):233–238.

Pleskac TJ, Yu S, Grunevski S, Liu T. Attention biases preferential choice by enhancing an option’s value. Journal of Experimental Psychology: General. 2023; 152(4):993.

Polania R, Woodford M, Ruff CC. Efficient coding of subjective value. Nature neuroscience. 2019; 22(1):134–142.

Rangel A, Hare T. Neural computations associated with goal-directed choice. Current opinion in neurobiology. 2010; 20(2):262–270.

Ratcliff R. A theory of memory retrieval. Psychological review. 1978; 85(2):59.

Ratcliff R, McKoon G. The diffusion decision model: theory and data for two-choice decision tasks. Neural computation. 2008; 20(4):873–922.

Ratcliff R, Voskuilen C, Teodorescu A. Modeling 2-alternative forced-choice tasks: Accounting for both magnitude and difference effects. Cognitive psychology. 2018; 103:1–22.

Roe RM, Busemeyer JR, Townsend JT. Multialternative decision field theory: A dynamic connectionst model of decision making. Psychological review. 2001; 108(2):370.

Roitman JD, Shadlen MN. Response of neurons in the lateral intraparietal area during a combined visual discrimination reaction time task. Journal of neuroscience. 2002; 22(21):9475–9489.

Rramani Q, Krajbich I, Enax L, Brustkern L, Weber B. Salient nutrition labels shift peoples’ attention to healthy foods and exert more influence on their choices. Nutrition Research. 2020; 80:106–116.

Sepulveda P, Usher M, Davies N, Benson AA, Ortoleva P, De Martino B. Visual attention modulates the integration of goal-relevant evidence and not value. Elife. 2020; 9:e60705.

Shadlen MN, Shohamy D. Decision making and sequential sampling from memory. Neuron. 2016; 90(5):927–939.

Shevlin BR, Smith SM, Hausfeld J, Krajbich I. High-value decisions are fast and accurate, inconsistent with diminishing value sensitivity. Proceedings of the National Academy of Sciences. 2022; 119(6):e2101508119.

Shimojo S, Simion C, Shimojo E, Scheier C. Gaze bias both reflects and influences preference. Nature neuroscience. 2003; 6(12):1317–1322.

Smith SM, Krajbich I. Attention and choice across domains. Journal of Experimental Psychology: General. 2018; 147(12):1810.

Smith SM, Krajbich I. Gaze amplifies value in decision making. Psychological science. 2019; 30(1):116–128.

Song M, Wang X, Zhang H, Li J. Proactive information sampling in value-based decision-making: Deciding when and where to saccade. Frontiers in human neuroscience. 2019; 13:35.

Steinemann NA, Stine GM, Trautmann EM, Zylberberg A, Wolpert DM, Shadlen MN. Direct observation of the neural computations underlying a single decision. bioRxiv. 2022; p. 2022–05.

Stine GM, Trautmann EM, Jeurissen D, Shadlen MN. A neural mechanism for terminating decisions. Neuron. 2023; 111(16):2601–2613.

Störmer VS, Alvarez GA. Attention alters perceived attractiveness. Psychological Science. 2016; 27(4):563–571.

Sullivan N, Hutcherson C, Harris A, Rangel A. Dietary self-control is related to the speed with which attributes of healthfulness and tastiness are processed. Psychological science. 2015; 26(2):122–134.

Summerfield C, Tsetsos K. Do humans make good decisions? Trends in cognitive sciences. 2015; 19(1):27–34.

Suzuki S, Cross L, O’Doherty JP. Elucidating the underlying components of food valuation in the human orbitofrontal cortex. Nature neuroscience. 2017; 20(12):1780–1786.

Tavares G, Perona P, Rangel A. The attentional drift diffusion model of simple perceptual decision-making. Frontiers in neuroscience. 2017; 11:468.

Thomas AW, Molter F, Krajbich I. Uncovering the computational mechanisms underlying many-alternative choice. Elife. 2021; 10:e57012.

Thomas AW, Molter F, Krajbich I, Heekeren HR, Mohr PN. Gaze bias differences capture individual choice behaviour. Nature Human Behaviour. 2019; 3(6):625–635.

Ting CC, Gluth S. High overall values mitigate gaze-related effects in perceptual and preferential choices. Journal of Experimental Psychology: General. 2025; .

Tolhurst D, Movshon JA, Thompson I. The dependence of response amplitude and variance of cat visual cortical neurones on stimulus contrast. Experimental brain research. 1981; 41:414–419.

Trueblood JS, Brown SD, Heathcote A. The multiattribute linear ballistic accumulator model of context effects in multialternative choice. Psychological review. 2014; 121(2):179.

Tversky A. Elimination by aspects: A theory of choice. Psychological review. 1972; 79(4):281.

Usher M, McClelland JL. The time course of perceptual choice: the leaky, competing accumulator model. Psychological review. 2001; 108(3):550.

Usher M, McClelland JL. Loss aversion and inhibition in dynamical models of multialternative choice. Psychological review. 2004; 111(3):757.

Vickers D. Decision processes in visual perception. Academic Press; 1979.

Wang XJ. Probabilistic decision making by slow reverberation in cortical circuits. Neuron. 2002; 36(5):955–968.

Westbrook A, Van Den Bosch R, Määttä J, Hofmans L, Papadopetraki D, Cools R, Frank M. Dopamine promotes cognitive effort by biasing the benefits versus costs of cognitive work. Science. 2020; 367(6484):1362–1366.

Yang X, Krajbich I. A dynamic computational model of gaze and choice in multi-attribute decisions. Psychological Review. 2023; 130(1):52.

Zajonc RB. Attitudinal effects of mere exposure. Journal of personality and social psychology. 1968; 9(2p2):1.

Zhu T. Accounting for the last-sampling bias in perceptual decision-making. Cognition. 2022; 223:105049.

Zylberberg A, Bakkour A, Shohamy D, Shadlen MN. Value construction through sequential sampling explains serial dependencies in decision making. Elife. 2024; 13:RP96997.

Zylberberg A, Fetsch CR, Shadlen MN. The influence of evidence volatility on choice, reaction time and confidence in a perceptual decision. Elife. 2016; 5:e17688.

